# Live-cell single-molecule fluorescence microscopy for protruding organelles reveals regulatory mechanisms of MYO7A-driven cargo transport in stereocilia of inner ear hair cells

**DOI:** 10.1101/2024.05.04.590649

**Authors:** Takushi Miyoshi, Harshad D. Vishwasrao, Inna A. Belyantseva, Mrudhula Sajeevadathan, Yasuko Ishibashi, Samuel M. Adadey, Narinobu Harada, Hari Shroff, Thomas B. Friedman

## Abstract

Stereocilia are unidirectional F-actin-based cylindrical protrusions on the apical surface of inner ear hair cells and function as biological mechanosensors of sound and acceleration. Development of functional stereocilia requires motor activities of unconventional myosins to transport proteins necessary for elongating the F-actin cores and to assemble the mechanoelectrical transduction (MET) channel complex. However, how each myosin localizes in stereocilia using the energy from ATP hydrolysis is only partially understood. In this study, we develop a methodology for live-cell single-molecule fluorescence microscopy of organelles protruding from the apical surface using a dual-view light-sheet microscope, diSPIM. We demonstrate that MYO7A, a component of the MET machinery, traffics as a dimer in stereocilia. Movements of MYO7A are restricted when scaffolded by the plasma membrane and F-actin as mediated by MYO7A’s interacting partners. Here, we discuss the technical details of our methodology and its future applications including analyses of cargo transportation in various organelles.

## 1. Introduction

Hearing loss affects approximately 20% of the population world-wide^1^ and is classified into either conductive or sensorineural hearing loss or a mixture of these two conditions^2^. Sensorineural hearing loss is usually caused by lesions in the inner ear originating from factors including pathogenic variants of genes, noise exposure, ototoxic drugs and aging. The exact etiology for hearing loss often remains unidentified in actual clinical practice^2^. In the inner ear, stereocilia of cochlear hair cells convert mechanical sound vibrations into electrochemical activities of neurons. Signals from hair cells are transmitted to the central nervous system through the afferent fibers of spiral ganglion neurons^3^. A similar mechanism is utilized by vestibular hair cells to detect acceleration including gravity^4^. Stereocilia are cylindrical F-actin protrusions formed on the apical surface of each hair cell. Degeneration of stereocilia often accompanies sensorineural hearing loss including those caused by aging and genetic pathogenic variants^5^. Despite recent advances in gene therapy such as for *OTOF*^6–8^, there are presently no clinically useful treatments to restore degenerated stereocilia or to regenerate lost hair cells in the cochlea^9^. Understanding the molecular events in stereocilia of live hair cells can be a basis to elucidate the pathophysiology of sensorineural hearing loss and to eventually formulate therapeutic strategies to restore normal hearing.

Single-molecule fluorescence microscopy is a powerful technique to localize molecules coupled with a fluorophore, such as GFP and organic fluorescent dyes^10^. One major application is real-time functional analyses of proteins and other molecules in live cells^11^ and on a coverslip between immobilized proteins and those in solution^12^. These techniques detect fluorescent molecules whose displacement, usually by diffusion, is smaller than the localization precision of an optical system^11^. Interactions between two fluorescently-labeled molecules can be detected more directly by Förster resonance energy transfer (FRET)^13^, optical tweezers^14^ and fluorescence correlation spectroscopy^15^. In addition, super-resolution images can be produced by serially localizing sparse subsets of fluorophores conjugated to imaging targets, for example, by STORM^16, 17^, PALM^18^ and PAINT^19, 20^ methodologies. These methods have been extended to three-dimensional tissue specimens by introducing a thin illumination plane and, if possible, minimizing out-of-focus interference using confocal microscopy^11, 21^. However, it has been challenging to apply single-molecule microscopy to organelles protruding from the apical plane, such as stereocilia, because most microscopes’ focal plane coincides with the plane of the imaging substrate. Volume scanning along the vertical axis is useful for these samples but has disadvantages in resolution, real-time acquisition and phototoxicity^22, 23^. One solution is to position a coverslip near the organelle of interest by placing samples upside-down on an inverted microscope^24^ or using a thin chamber^25^ although these techniques risk damaging the organelles and are not suitable for drug treatments. We solved this problem by employing a dual-view inverted selective plane illumination microscope (diSPIM)^26^ and established a workflow of single-molecule microscopy with a more flexible spatial arrangement. We first imaged the dynamics of MYO7A, one of the unconventional myosins essential for normal hearing and vision^27, 28^.

Development of functional stereocilia and their maintenance for life-long hearing require regulated motor activity of the wild-type unconventional myosins MYO3A, MYO6, MYO7A and MYO15A^29, 30^. Pathogenic variants of these myosin genes are associated with human hereditary nonsyndromic hearing loss^27^. Variants of *MYO7A* are also associated with Usher syndrome type 1 characterized by congenital hearing loss, vestibular dysfunction and progressive retinal degeneration leading to blindness in the second to third decade of life^28^. During the development of hair cells, microvilli-like F-actin protrusions are formed on the apical surface of immature hair cells, of which a majority are “pruned” and only a small subset of microvilli grow thicker and taller and develop cross-linked paracrystal-like F-actin bundles in them. These maturing stereocilia are organized in rows of increasing height. All rows of stereocilia except the tallest are equipped with mechanoelectrical transduction (MET) channels at the distal ends. Gating of MET channels is mediated by tip-links physically connected to a spot on the side of adjacent longer stereocilia^31^, which is referred to as the upper tip-link density (UTLD) because of the high electron scattering in transmission electron micrographs^3^ (see Fig. 6a). Opening of MET channels allows K^+^ and Ca^2+^ from the endolymph to enter stereocilia and depolarizes the plasma membrane, which leads to release of glutamate from the basal surface of the hair cell^32–34^. Unconventional myosins are involved in these processes by transporting and anchoring specific proteins and phospholipids as “cargo” using their unique tail domains as binding sites^35^.

A number of proteins have been identified as “cargo” of unconventional myosins in stereocilia. For example, MYO15A transport factors to elongate F-actin cores including WHRN (whirlin) and EPS8 (epidermal growth factor receptor kinase substrate 8)^36, 37^. The motor domain of MYO15A itself is also reported to nucleate actin monomers^38^. MYO3A can interact with ESPN (espin) isoform 1 and ESPNL (espin-like), both of which are crucial for elongating F-actin protrusions^39, 40^, although localization of these proteins in stereocilia may not completely depend on the motor activities of class-III myosins^41^. MYO7A localizes in the UTLD along with two scaffolding proteins, SANS and Harmonin, and tethers the tip-link consisting of CDH23 and PCDH15 to the F-actin core^42–44^. It is obvious that the motor activities of these myosins are necessary for localizing their cargo because mice with missense mutations disabling the motor domain of MYO7A and MYO15 have profound hearing loss^45, 46^. However, less is understood about how these unconventional myosins traffic and transport cargo in stereocilia using energy from ATP hydrolysis. Mutant mouse models may not always be suitable to approach this question because stereocilia are often severely deformed in mice with impaired activity of these myosins, which likely disrupts cargo transport^45^.

In this study, we develop the methodology for live-cell single-molecule microscopy applicable to organelles protruding from the apical surface of tissue using a dual-view light-sheet microscope, diSPIM^26^, and bright fluorescent dyes (Fig. 1). Here, we experimentally address the unresolved question of how MYO7A molecules can transport components of the tip-link complex (Figs. 2–6). MYO10, an unconventional myosin crucial for filopodia formation, was used as a positive control. We found that MYO7A shows directional, processive movements toward stereocilia tips when the motor domain is exposed by disabling its tail-mediated motor inhibition^47, 48^. We further compared movements of MYO7A motor domains in stereocilia among (1) homodimers, (2) monomers anchored to the plasma membrane and (3) monomers tethered to F-actin at the C-terminus, all of which are possible through interacting partners of MYO7A. Among these conditions, only homodimers showed processive movements suggesting that MYO7A moves as a dimer in a stereocilium. The knowledge of how myosin-driven cargo transport occurs in stereocilia will be applicable to other F-actin protrusions, microvilli and filopodia, where diffusion of proteins are limited in a tightly-packed F-actin core and in an envelope of the plasma membrane^49^. Here, we introduce the technical details and methodology of our single-molecule microscopy of live hair cell stereocilia and its general application to three-dimensional specimens including analyses of cargo transportation mechanisms in various organelles.

**Fig. 1:**
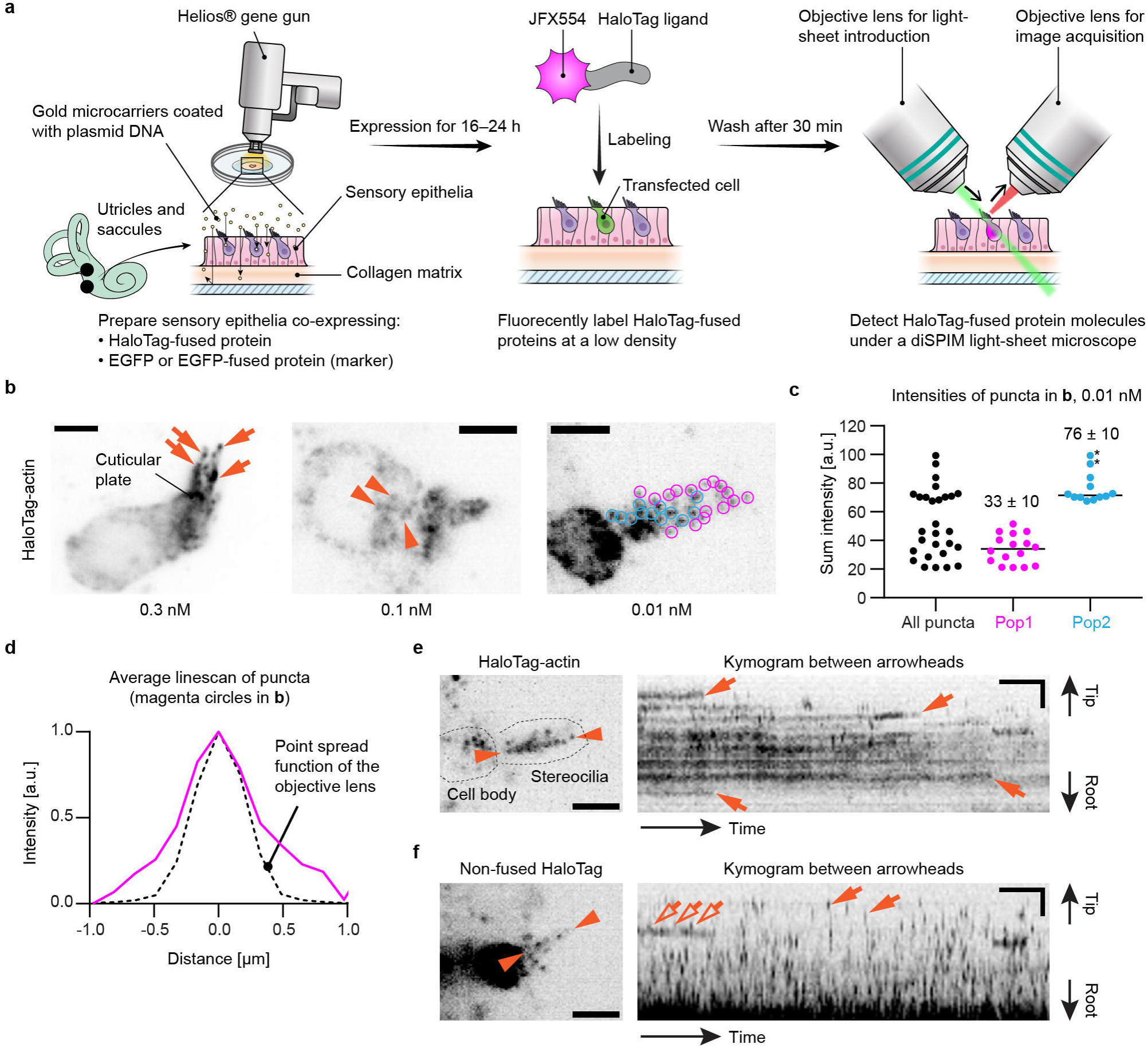
Development of single-molecule microscopy in live hair cells. **a,** Illustration showing our workflow for single-molecule microscopy. **b,** Optimization of labeling density using vestibular hair cells expressing HaloTag-actin. JFX554-conjugated HaloTag-ligands applied at different concentrations. At 0.3 nM or above, JFX554-ligands distribute throughout the cell and accumulated at stereocilia tips (arrows) and in the cuticular plate. Fluorescent puncta begin to appear at 0.1 nM in the cell body (arrowheads) and becomes distinguishable in stereocilia around 0.01 nM (indicated by colored circles for quantification in **c**). Maximum projections of volume scans are shown. Exposure, 100 ms per plane at 0.2 kW/cm^2^. Bars, 5 µm. **c**, Intensities of fluorescent puncta in **b**, 0.01 nM JFX554-ligand. The sum intensity of each punctum is calculated by adding all pixel values encompassing the punctum and then subtracting the background intensity. There are at least two populations of fluorescent puncta, Pop1 and Pop2, indicated by magenta and cyan circles in **b**, respectively. The average intensity of Pop2 is 76 ± 10 (n = 12; mean ± standard deviation) and approximately twice as large as that of Pop1 (34 ± 10; n = 16) indicating puncta in Pop1 and Pop2 are emitted from one and two fluorophores, respectively. Puncta of high intensity in Pop2 (asterisks) may be emitted from more than two fluorophores. Pixel values are calculated using an average projection of the volume scan. **d**, Comparison between the average line-scan of fluorescent puncta (magenta circles in **b**, 0.01 nM) and the point spread function of the objective lens calculated using PSF Generator (https://bigwww.epfl.ch/algorithms/psfgenerator/). Similarity between both intensity curves suggests that these puncta are emitted from a point source. **e and f**, Representative kymograms of controls, HaloTag-actin (**e**) and non-fused HaloTag (**f**) labeled with 0.01 nM and 0.1 nM JFX554-ligands, respectively. Single-plane images are acquired every 1 s for comparison with MYO7A movements. Kymograms are generated from the line scans between arrowheads. Most HaloTag-actin molecules stay in the same place and disappear suddenly due to photobleaching or transition to the dark state (**e**, arrows), which also suggests that these puncta are emitted from single fluorophores. Most non-fused HaloTag molecules disappear after one frame (**f**, arrows) due to diffusion except for a few molecules likely stuck in stereocilia (**f**, open arrows). Imaging conditions are similar to **b**. Bars, 20 s and 2 µm.

**Fig. 2:**
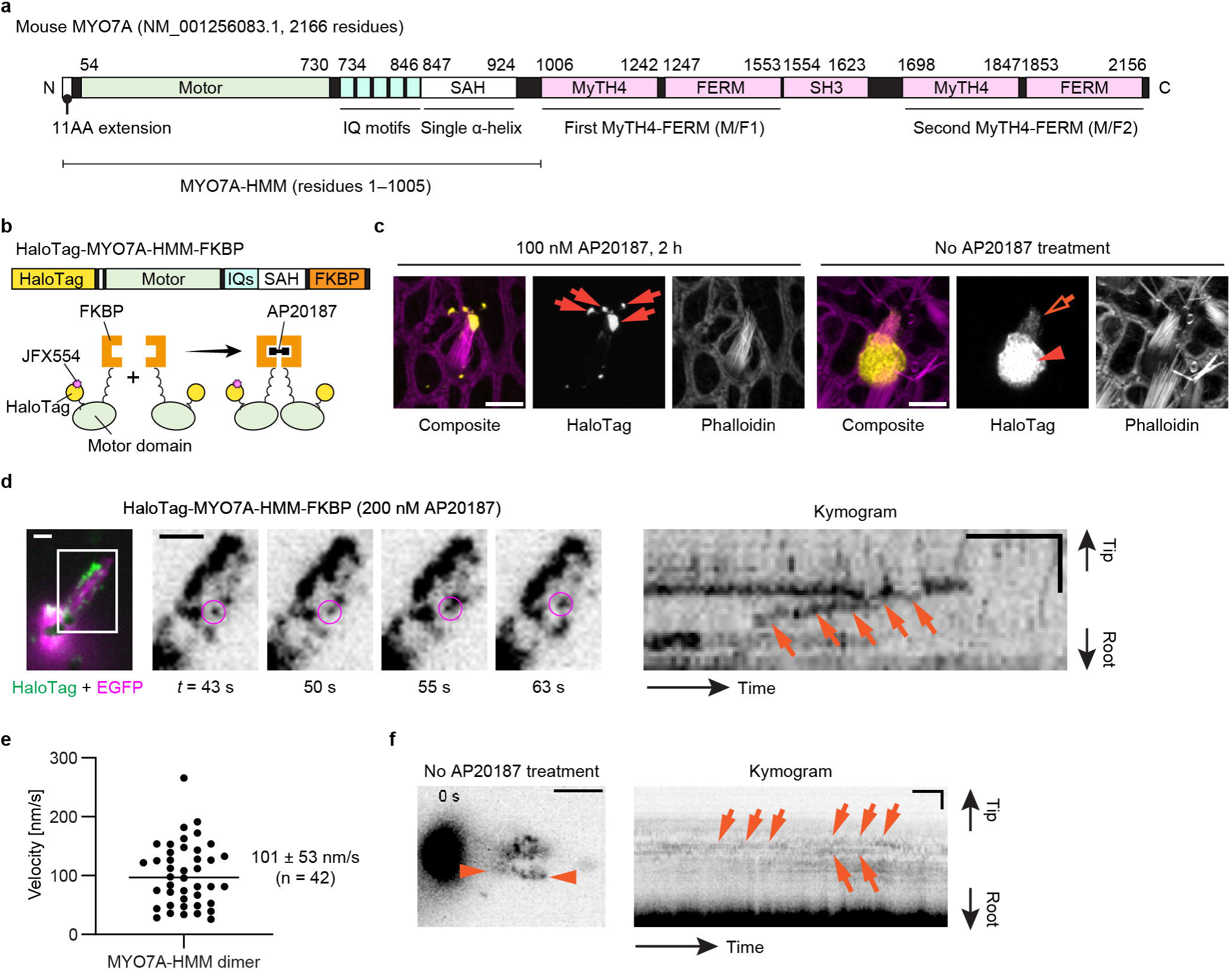
Imaging of MYO7A-HMM dimers directionally moving in stereocilia. **a,** Domain structures of mouse MYO7A (NM_001256083.1) and the heavy-meromyosin-like fragment (MYO7A-HMM) used in this study. **b,** Schemes showing the structure of HaloTag-MYO7A-HMM-FKBP and conditional dimerization under AP20187 treatment. Note that only a small portion of HaloTag-fused protein is labeled in our single-molecule microscopy. **c,** AP20187-dependent accumulation of HaloTag-MYO7A-HMM-FKBP at stereocilia tips. Vestibular hair cells (P2) expressing HaloTag-MYO7A-HMM-FKBP are incubated with or without 100 nM AP20187 for 2 h. Samples are fixed and stained by 200 nM JFX554-conjugated HaloTag ligands (yellow) and Alexa405-phalloidin (magenta) for confocal microscopy. HaloTag-MYO7A-HMM-FKBP accumulates at stereocilia tips and forms large protein blobs in cells treated with AP20187 (arrows), while HaloTag-MYO7A-HMM-FKBP without AP20187 treatment localizes diffusely in the cuticular plate (arrowhead) and stereocilia (open arrow). These localization patterns are consistent with AP20187-dependent directional movements of HaloTag-MYO7A-HMM-FKBP toward stereocilia tips. Bars, 5 µm. **d,** Processive movements of MYO7A-HMM dimers in stereocilia under 200 nM AP20187 treatment. Time-lapse images (black-and-white) of magnified white rectangle area in the pseudo-colored image (HaloTag - green, EGFP - magenta) show a directionally moving molecule in stereocilia (magenta circles). The continuous trajectory on the kymogram indicates its processive movement (arrows). JFX554, 0.3 nM. Single-plane time-lapse, every 1 s. Bars, 5 µm (time-lapse images); 2 µm and 20 s (kymogram). **e,** Velocities of MYO7A-HMM dimers moving in stereocilia. The average is 101 ± 53 nm/s (n = 42, mean ± standard deviation). **f,** Behavior of HaloTag-MYO7A-HMM-FKBP molecules in untreated cells. Trajectories in the kymogram (between arrowheads) are parallel to the time axis (arrows) indicating that HaloTag-MYO7A-HMM-FKBP molecules remain at the same position. Imaging conditions and scale bars are the same as in **d**.

## 2. Results

### 2.1. Single-molecule microscopy in stereocilia of live hair cells

The workflow for our single-molecule microscopy approach was developed using explant cultures of mouse utricles and saccules, hereafter referred to as “vestibular sensory epithelia”, harvested from postnatal day (P) 2 to 5 (Fig. 1a). Stereocilia of utricles and saccules are suitable for single-molecule microscopy because they are straight and can be as long as 10 µm^50^. We employed a dual-view inverted selective plane illumination microscope (diSPIM)^26^ in order to image stereocilia protruding upward from the apical surface of hair cells as we previously reported for stereocilia of fixed hair cells^12^. Using a Helios® gene-gun^51^, explant cultures of vestibular sensory epithelia were co-transfected with expression plasmids encoding a HaloTag-fused protein of interest and an EGFP (or EGFP-fused protein) to function as a transfection marker. Transfected vestibular sensory epithelia were maintained in DMEM/F12 culture media (37°C, 5% CO_2_) and allowed to express these proteins for 16–24 hours. HaloTag-fused protein was fluorescently labeled at a low density with JFX554-conjugated HaloTag ligands^52^ and imaged with single-molecule microscopy using a diSPIM^26^ illuminating a 561-nm laser. To image single protein molecules, we took advantage of our previously reported methodology of multiplexed super-resolution microscopy, in which we also detected single molecules of fluorescently labeled imaging probes with a diSPIM^12^.

The concentration of JFX554-conjugated HaloTag ligands was optimized using vestibular hair cells expressing HaloTag-fused human β-actin (HaloTag-actin) (Fig. 1b). With ligands applied at 0.3 nM or higher, the entire cell was labeled although more densely at stereocilia tips and in the cuticular plate (Fig. 1b, arrows), similarly to hair cells expressing EGFP-actin^24^. Fluorescent puncta of single HaloTag-actin molecules appeared in the cell body at concentrations of 0.1 nM (Fig. 1b, arrowheads) and in stereocilia at ligand concentrations of 0.03 nM or lower (Fig. 1b, circles, image of 0.01 nM shown). The optimal concentration was slightly different between cells depending on the amount of expressed HaloTag-actin. With dyes at 3 nM or above, unreacted fluorescent dyes were not completely washed away and remained in the tissue (image not shown). Thus, we considered that fluorescent ligands should be applied below 3 nM and be optimized depending on the expression level of HaloTag-fused protein. We also confirmed that single protein molecules are visualized by calculating the summed fluorescence intensity for each fluorescent punctum (Fig. 1c) using the cell in Fig. 1b, 0.01nM. Fluorescent puncta were able to be classified into two populations in this cell (indicated by magenta and cyan circles in Fig. 1b, 0.01 nM). The average intensity of the Pop2 population was twice as large as that of the Pop1 population in Fig. 1c, indicating that fluorescent puncta in Pop1 originated from one fluorophore and that the average intensity of Pop1 corresponds to the quantal intensity. Concordantly, the average line scan of puncta in Pop1 was consistent with the point spread function of the objective lens (Fig. 1d).

Time-lapse images of HaloTag-actin and non-fused HaloTag were acquired to evaluate how proteins are visualized when stably bound to the F-actin core and diffusing in stereocilia, respectively (Fig. 1, e and f). Single-plane time-lapse images were acquired every 1 second (s) to compare with subsequent MYO7A imaging (Figs. 2–6). In the kymogram and movie, most HaloTag-actin molecules showed trajectories parallel to the time axis (X-axis) and disappeared suddenly due to photobleaching or transition to the dark state (Fig. 1e, arrows and Movie S1). This behavior is consistent with single fluorophores staying in the same position. Consistent with diffusion, almost all non-fused HaloTag molecules showed trajectories no longer than one frame without staying in the same position (Fig. 1f, arrows and Movie S2) except for a few molecules stuck in stereocilia (Fig. 1f, open arrows). From these data, we concluded that fluorescent puncta in the time-lapse images reflect the dynamics of single protein molecules.

### 2.2. Visualization of directional movement of MYO7A dimers in stereocilia

After establishing the workflow for single-molecule microscopy, we developed imaging conditions suitable for detecting directional movements of HaloTag-fused MYO7A molecules in stereocilia (Fig. 2). We employed the heavy meromyosin-like fragment of mouse MYO7A (MYO7A-HMM) for this purpose (Fig. 2a). MYO7A-HMM is designed to be similar in domain composition to the heavy meromyosin (HMM)^53^, a protein fragment obtained by myosin II trypsinization and consisting of the motor and neck domains necessary for a power stroke on F-actin^54, 55^. MYO7A-HMM dimers show directional movements in filopodia and microvilli^53, 56^ and are expected to be a useful benchmark in stereocilia. In addition, MYO7A-HMM can be conditionally dimerized in live cells by fusing the p.F36V substitution mutation of FK506 binding protein 12 (FKBP) to the C-terminus and by adding a FK506-derived bivalent ligand, AP20187, to the culture medium^57^. This chemically inducible dimerization can “turn on” trafficking of MYO7A-HMM when the cells are ready for imaging. Thus, we constructed an expression vector for HaloTag-MYO7A-HMM-FKBP, which has a HaloTag for fluorescent labeling at the N-terminus and the FKBP for conditional dimerization at the C-terminus (Fig. 2b). HaloTag-MYO7A-HMM-FKBP expressed in vestibular hair cells formed large protein blobs at stereocilia tips only when AP20187 was added to the culture medium indicating that MYO7A-HMM dimers move toward the barbed ends of unidirectional F-actin bundles in stereocilia cores (Fig. 2c).

Single HaloTag-MYO7A-HMM-FKBP molecules were successfully detected using the imaging condition established with HaloTag-actin but at a slightly higher concentration of JFX554-ligands, 0.3–0.6 nM (Fig. 2d, Movie S3). MYO7A required a higher concentration of JFX554-ligand than β-actin likely reflecting the different expression levels of these two proteins. Time-lapse images after the AP20187 treatment visualized HaloTag-MYO7A-HMM-FKBP molecules moving directionally toward stereocilia tips (Fig. 2d, magenta circles). Kymograms showed continuous trajectories consistent with processive movements of MYO7A-HMM dimers (Fig. 2d, arrows). The velocity of movement was different between dimers (Fig. S1; representative kymograms) although there are no distinct populations of “slow” and “rapid” movements (Fig. 2e). The average velocity of movements was 101 ± 53 nm/s (n = 42; mean ± standard deviation), which is 10-fold faster than the movements of human recombinant MYO7A-HMM dimers on permeabilized filopodia (9.5 ± 0.4 nm/s)^58^. As we discuss later, this difference can be partially attributed to the temperature (37°C in our study vs. 25°C in the previous study) considering that similar difference was observed for single-molecule microscopy of MYO10 in live-cell filopodia (578 ± 174 nm/s at 25°C vs. 840 ± 210 nm/s at 37°C)^59^. HaloTag-MYO7A-HMM-FKBP did not show directional movements without AP20187 (Fig. 2f, Movie S4).

### 2.3. Constitutively active MYO7A mutants move directionally in stereocilia

Using the imaging condition established with MYO7A-HMM dimers, we tested the hypothesis that MYO7A traffics as dimers (or oligomers) in stereocilia (Fig. 3). However, HaloTag-fused full-length MYO7A did not show directional movements at a detectable frequency in stereocilia of vestibular hair cells (image not shown). We considered the possibility that full-length MYO7A takes a backfolded autoinhibitory conformation between the tail and motor domains^47, 48^. These studies also indicate that this autoinhibitory interaction is mediated by the “RGSK” motif in the second MyTH4-FERM domain (M/F2) and can be disabled by substituting arginine and lysine in this motif with alanine residues. Thus, we evaluated movements of two HaloTag-fused MYO7A mutants, HaloTag-MYO7A-RK/AA and HaloTag-MYO7A-ΔSH3-ΔM/F2, which disable the tail-mediated autoinhibition by missense mutations of RK to AA residues and by a deletion of SH3 and M/F2 domains, respectively (Fig. 3a). While wild-type MYO7A diffusely distributed in stereocilia (Fig. 3b, arrowhead in Wild-type), these MYO7A mutants accumulated at stereocilia tips in a few transfected hair cells (Fig. 3b, arrows in RK/AA and ΔSH3-ΔM/F2). We speculate that accumulation at stereocilia tips may occur only when sufficient amounts of HaloTag-fused MYO7A and its interaction partners are present in the cell.

**Fig. 3:**
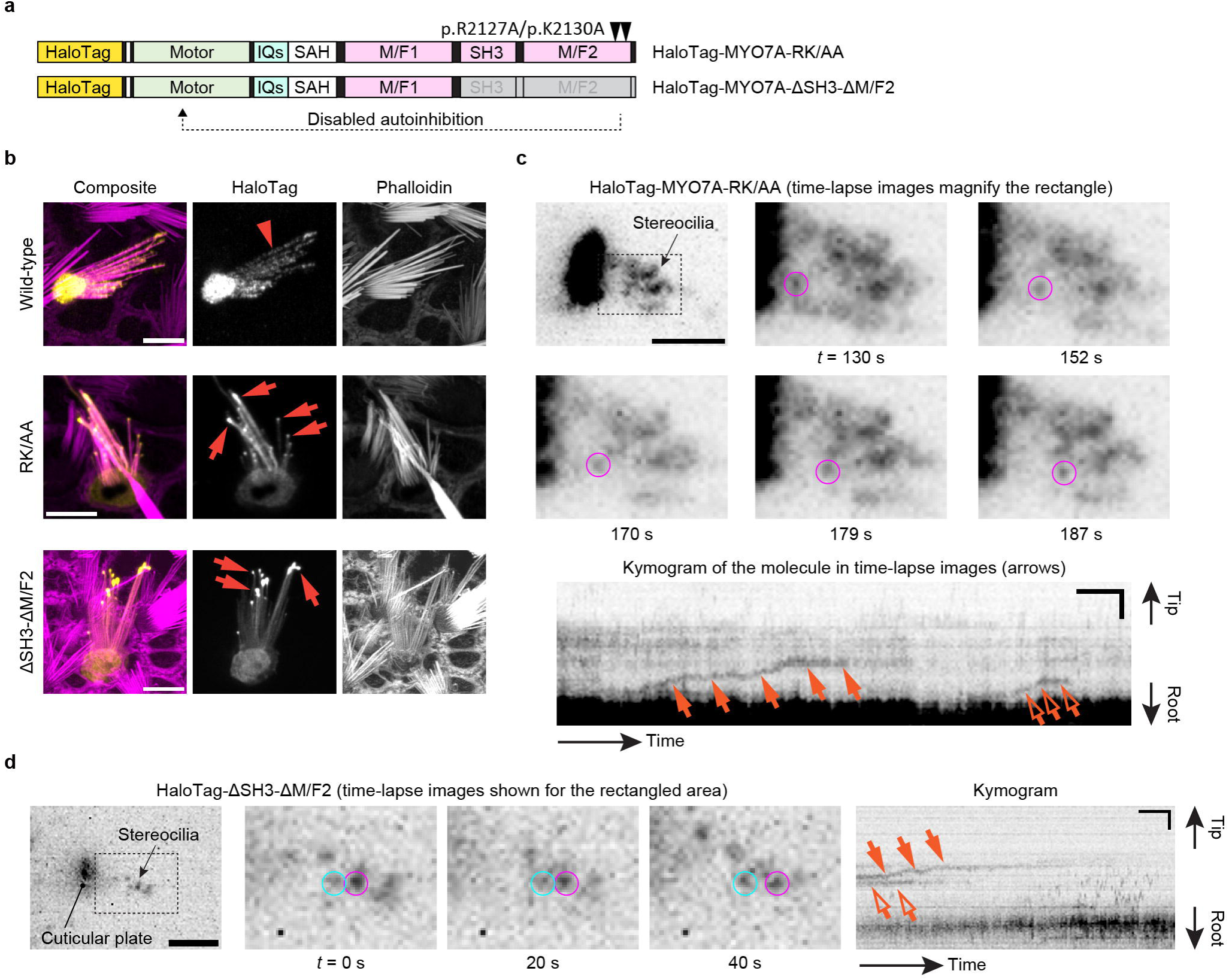
Directional movements of constitutively active MYO7A mutants. **a,** Structures of two constitutively active MYO7A mutants fused with HaloTag at the N-terminus. HaloTag-MYO7A-RK/AA has two missense mutations, p.R2127A and p.K2130A, that we inserted in the second MyTH4-FERM domain (M/F2) referring to the study using human MYO7A^111^. HaloTag-MYO7A-ΔSH3-ΔM/F2 has a truncated tail. These mutations were introduced to remove the tail-mediated autoinhibition of the motor domain. **b,** Accumulation of constitutively active MYO7A mutants at stereocilia tips. Vestibular hair cells (P2–5) expressing HaloTag-fused wild-type MYO7A and the two mutants described in **a** are subjected to confocal microscopy. Wild-type MYO7A diffusely distributes in stereocilia (arrowhead), while MYO7A mutants accumulate at stereocilia tips (arrows) in some cells suggesting directional movements of these mutants toward stereocilia tips. Bars, 5 µm. **c**, Single-molecule microscopy of HaloTag-MYO7A-RK/AA. Time-lapse images show a molecule directionally moving in stereocilia (magenta circles). Kymogram illustrates the processive movement of this molecule (arrows) and another directionally moving molecule (open arrow). JFX554, 0.3 nM. Single-plane time-lapse, every 1 s. Bars, 5 µm (time-lapse images); 2 µm and 20 s (kymogram). **d**, Single-molecule microscopy of HaloTag-MYO7A-ΔSH3-ΔM/F2. Time-lapse images and kymogram show processive and directional movement of a molecule (magenta circles and arrows). A static molecule is indicated for comparison (cyan circles and open arrows). Imaging conditions and scale bars are the same as in **c**.

Under single-molecule imaging conditions, a few HaloTag-MYO7A-RK/AA molecules moved toward stereocilia tips (Fig. 3c, Movie S5). Movements of MYO7A-RK/AA were directional and processive resembling the movements of MYO7A-HMM dimers. This result suggests that MYO7A can dimerize spontaneously on the F-actin cores of stereocilia when the motor domain is exposed, for example, by cargo bound to the tail^47, 60, 61^. Similar processive movements were observed in cells expressing HaloTag-MYO7A-ΔSH3-ΔM/F2 (Fig. 3d, Movie S6) indicating that MYO7A can dimerize using motifs in the neck or in the first MyTH4-FERM domain (M/F1).

### 2.4. Distinct behavior of MYO7A and MYO10 anchored to the plasma membrane

Our data indicate that MYO7A can traffic in stereocilia as a dimer or perhaps an oligomer. Another possibility is that MYO7A is anchored to the plasma membrane of stereocilia and showed directional movements on the adjacent F-actin core (Fig. 4). Previous studies show that MYO7A can interact with CDH23 via two scaffolding proteins in the tip-link complex, SANS and Harmonin^62–64^, and directly with PCDH15^65^ (see Fig. 6b). In addition, anchoring to the plasma membrane can induce directional movements of MYO10 in filopodia^66^. Thus, we tested if MYO7A-HMM can show directional movements when tethered to the plasma membrane. Anchoring of MYO7A-HMM to the plasma membrane was achieved using a small transmembrane motif from the human Interleukin 2 receptor alpha chain (IL2Rα)^67^ (Fig. 4a). IL2Rα and MYO7A-HMM were conditionally heterodimerized by fusing FKBP and FKBP-Rapamycin binding protein (FRB) to the C-terminus of each protein and supplementing the culture medium with a Rapamycin analog, AP21987^68^. Bovine MYO10 lacking the entire tail and the coiled-coil domain for anti-parallel dimerization (MYO10-MD) in the neck^66^ was used as a positive control. Fluorescence confocal microscopy shows that HaloTag-MYO7A-HMM-FRB is successfully anchored to the plasma membrane after AP21987 treatment (Fig. 4b). The positive control, HaloTag-MYO10-MD-FRB, accumulated weakly at stereocilia tips in a few cells without AP21987 in the medium (Fig. 4c, arrowhead) and more densely at stereocilia tips after the addition of AP21987 (Fig. 4c, arrows) as previously reported for filopodia^66^. Excess IL2Rα-EGFP-FKBP sometimes accumulated in vesicles but did not cause apparent damage to stereocilia (Fig. 4b, open arrowheads).

**Fig. 4:**
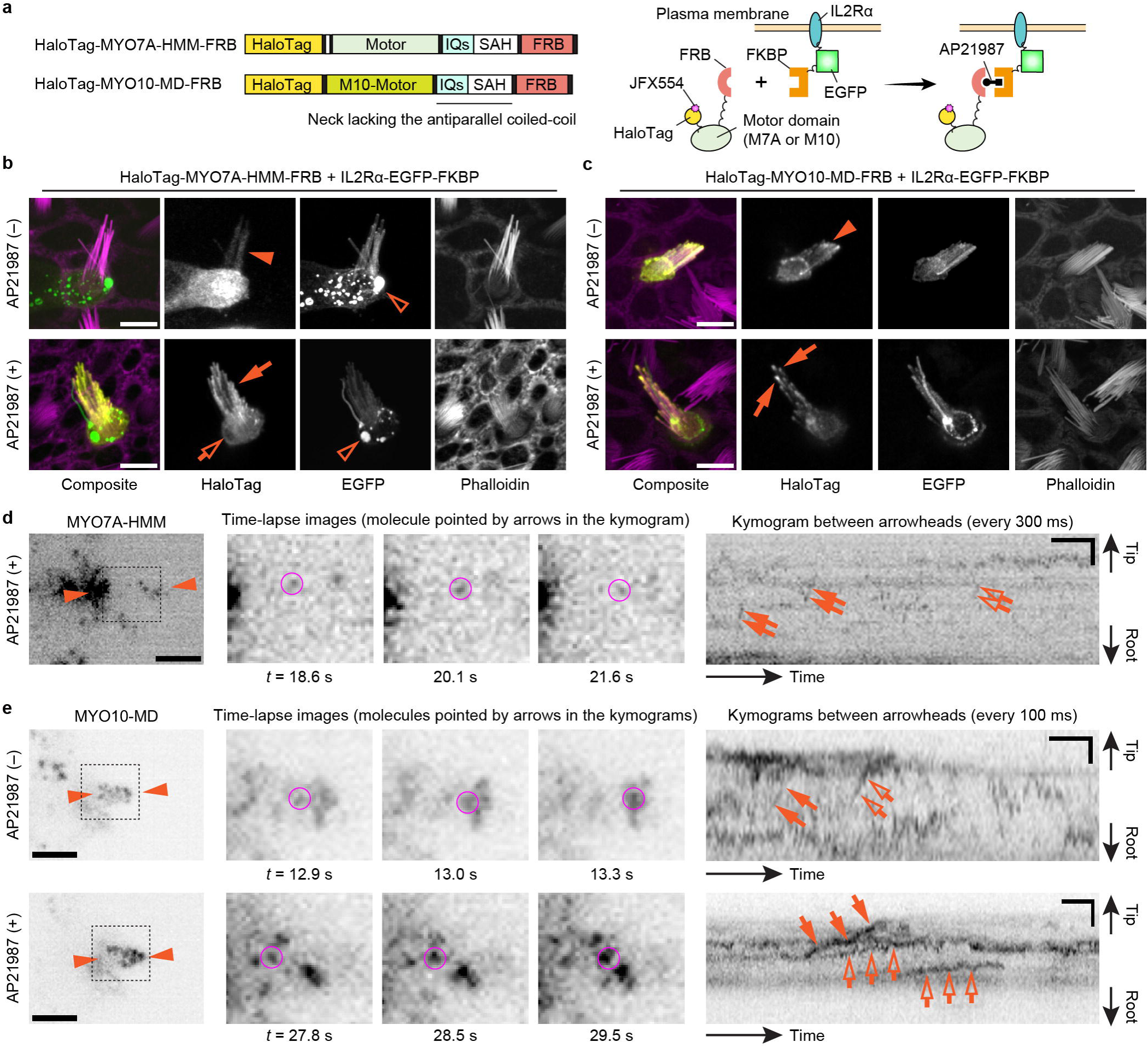
Movements of membrane-anchored MYO7A-HMM and MYO10-MD. **a,** Domain structures of HaloTag-MYO7A-HMM-FRB and HaloTag-MYO10-MD-FRB and a scheme showing membrane anchoring by IL2Rα-EGFP-FKBP (human Interleukin 2 receptor alpha chain fused with EGFP and FKBP). Membrane anchoring is mediated by conditional heterodimerization between FRB and FKBP under the AP21987 treatment. MYO10-MD is a control myosin motor head that can utilize the plasma membrane as a scaffold for directional movements in filopodia^66^. **b**, Confocal images showing AP21987-dependent membrane anchoring of MYO7A-HMM. Vestibular hair cells (P2) co-expressing HaloTag-MYO7A-HMM-FRB and IL2Rα-EGFP-FKBP are incubated with or without 500 nM AP21987 for 2 h. HaloTag-MYO7A-HMM-FRB distributes diffusely in stereocilia of untreated cells (arrowhead) but accumulates along the plasma membrane of stereocilia (arrow) and at the edge of the cuticular plate (open arrow) in AP21987-treated cells. Excess IL2Rα-EGFP-FKBP sometimes accumulates in vesicles within the cuticular plate without apparent damage to stereocilia (open arrowheads). Bars, 5 µm. **c**, Confocal images showing AP21987-dependent membrane anchoring of MYO10-MD and its localization changes. HaloTag-MYO10-MD-FRB accumulates at stereocilia tips weakly in a few untreated cells (arrowhead) suggesting that a small number of MYO10-MD molecules can move directionally without a scaffold. Increased accumulation of HaloTag-MYO10-MD-FRB at stereocilia tips in AP21987-treated cells (arrows) indicates directional movements enhanced by membrane anchoring. These localizations in stereocilia are consistent with a previous study of MYO10-MD in filopodia^66^. Bars, 5 µm. **d**, Single-molecule microscopy of membrane-anchored MYO7A-HMM. Staircase-like trajectories in kymogram indicate step-wise movements of MYO7A-HMM (arrows and open arrows). The molecule indicated by arrows is shown in time-lapse images (magenta circles). AP21987, 500 nM. Single-plane time-lapse, every 300 ms. Bars, 5 µm (cell image); 6 s and 2 µm (kymogram). **e,** Single-molecule microscopy of MYO10-MD. Before the AP21987 treatment, a small number of MYO10-MD molecules show rapid directional movements toward stereocilia tips (arrows and open arrows in the upper kymogram). After the AP21987 treatment, MYO10-MD molecules show slow directional movements (arrows and open arrows in the lower kymogram). Molecules indicated by arrows are shown in time-lapse images (magenta circles). AP21987, 500 nM. Single-plane images, every 100 ms. Bars, 5 µm (cell images); 2 s and 2 µm (kymograms).

Single-molecule microscopy showed that MYO7A-HMM can move in a stereocilium using the plasma membrane as a scaffold although the movements were restricted and different from those of dimers (Fig. 2d and Fig. 4d). In contrast, MYO10-MD moved efficiently toward stereocilia tips when anchored to the plasma membrane (Fig. 4e). In cells expressing HaloTag-MYO7A-HMM-FRB and IL2Rα-EGFP-FKBP, only a small number of MYO7A-HMM molecules moved directionally toward stereocilia tips after AP21987 treatment (Fig. 4d, Movie S7). Movements of membrane-anchored MYO7A-HMM were intermittent in a kymogram as if MYO7A-HMM moves toward stereocilia tips only when they dissociate from F-actin (Fig. 4d, arrows and open arrows). As indicated by the weak accumulation in fluorescent histochemistry (Fig. 4c, arrowhead), a small number of MYO10-MD molecules moved directionally even in untreated cells (Fig. 4e, arrows and open arrows in the upper kymogram, Movie S8). After AP21987 treatment, MYO10-MD anchored to the plasma membrane showed processive movements toward stereocilia tips (Fig. 4e, arrows and open arrows in the lower kymogram, Movie S9; representative kymograms also in Fig. S2b). The average velocity of MYO10-MD was 2.01 ± 0.37 μm/s (n = 12) before treatment and 0.72 ± 0.34 µm/s (n = 23) after treatment (Fig. S2a) indicating that movements of MYO10-MD were affected by membrane anchoring. Retrograde movements were also observed for MYO10-MD (Fig. S2b, arrowheads). Movements of membrane-anchored MYO7A-HMM were different from MYO7A-HMM dimers or constitutively active MYO7A mutants. Directional movements of MYO10-MD indicate that the restricted movements of MYO7A-HMM are derived from the kinetic differences between the motor domains of MYO7A and MYO10.

### 2.5. Step-wise movements of MYO7A and MYO10 when tethered to F-actin

We tested an additional scenario where F-actin functions as a scaffold for MYO7A to move in a stereocilium (Fig. 5). In the UTLD of stereocilia, a class of Harmonin isoforms that contain the Proline, Serine and Threonine-rich (PST) domain (collectively referred to as Harmonin b) connect the tip-link complex to the F-actin core including MYO7A^62^. Thus, we conditionally tethered HaloTag-MYO7A-HMM-FKBP or HaloTag-MYO10-MD-FKBP to F-actin under AP21987 treatment using the PST domain of Harmonin b (residues 296–728 of NM_01163733) fused with FRB and EGFP at the N- and C-termini (FRB-PST-EGFP) (Fig. 5a). Before AP21987 treatment, both HaloTag-MYO7A-HMM-FKBP and HaloTag-MYO10-MD-FKBP diffusely distributed in stereocilia (Fig. 5, b and c, arrowheads). FRB-PST-EGFP also distributed diffusely in stereocilia and weakly accumulated at stereocilia tips (Fig. 5, b and c, open arrowheads). After AP21987 treatment, HaloTag-MYO7A-HMM-FKBP co-localized with FRB-PST-EGFP at the tips (Fig. 5b, arrows). This result indicates that HaloTag-MYO7A-HMM-FKBP successfully heterodimerized with FRB-PST-EGFP. HaloTag-MYO10-MD-FKBP formed protein blobs at stereocilia tips with FRB-PST-EGFP after AP21987 treatment (Fig. 5c, arrows and open arrows) suggesting that MYO10-MD can move directionally and accumulate at stereocilia tips using F-actin as a scaffold.

**Fig. 5:**
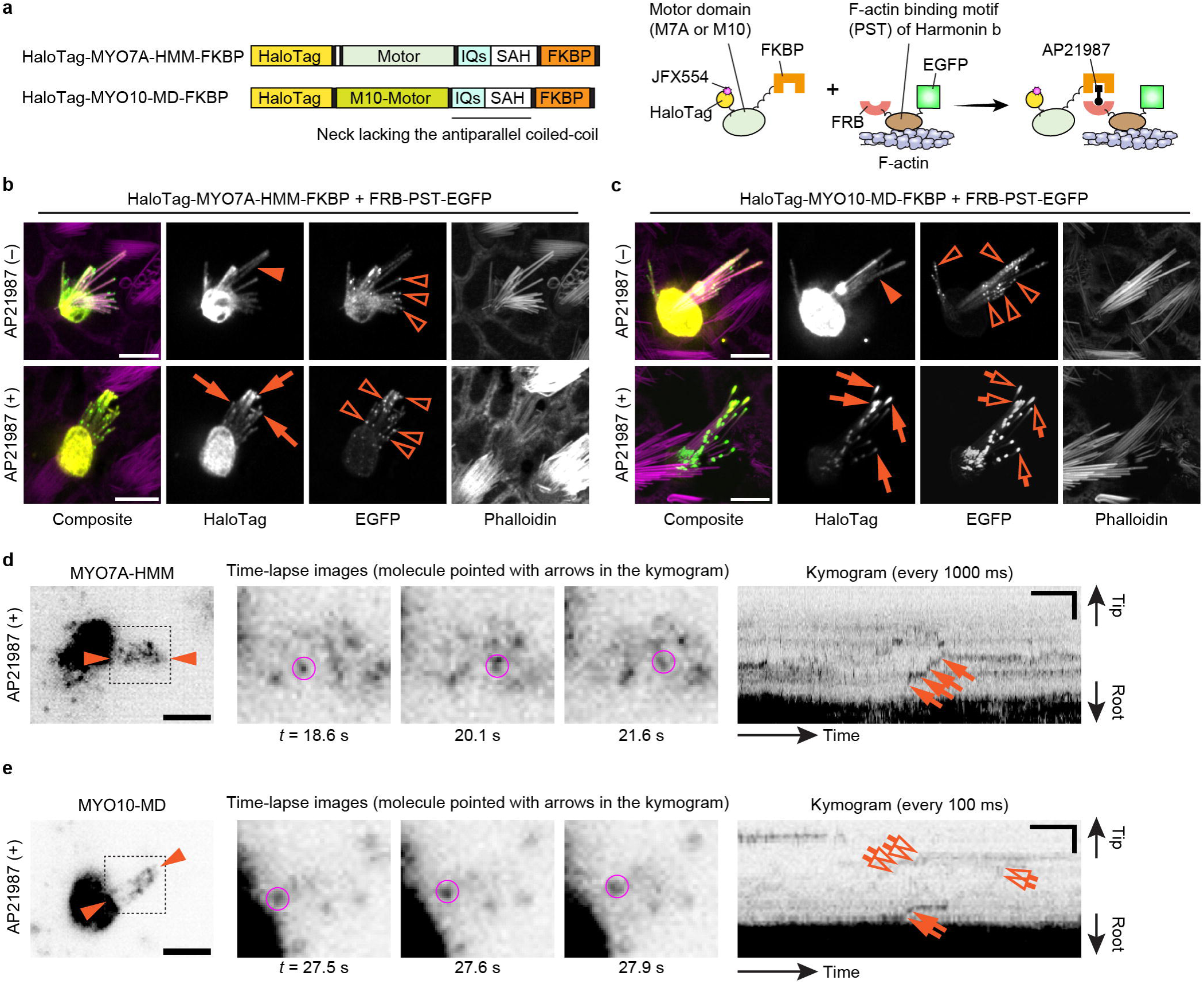
Step-wise movements of MYO7A-HMM and MYO10-MD tethered to F-actin. **a,** Scheme showing domain structures of HaloTag-MYO7A-HMM-FKBP and HaloTag-MYO10-MD-FKBP and AP21987-induced tethering to F-actin via FRB-PST-EGFP, the PST domain of mouse Harmonin b (residues 296–728 of NM_01163733) fused with FRB and EGFP at the N- and C-termini, respectively. **b**, Confocal images consistent with AP21987-dependent binding between MYO7A-HMM and PST. Vestibular hair cells (P2) co-expressing HaloTag-MYO7A-HMM-FKBP and FRB-PST-EGFP are incubated with or without 500 nM AP21987 for 2 h. FRB-PST-EGFP weakly accumulates at stereocilia tips (open arrowheads). HaloTag-MYO7A-HMM-FKBP distributes diffusely in untreated cells (arrowhead) and co-localizes with FRB-PST-EGFP only in cells treated with AP21987 (arrows). Bars, 5 µm. **c**, Confocal images consistent with AP21987-dependent binding between MYO10-MD and PST and a change in MYO10-MD localization. FRB-PST-EGFP weakly accumulates at stereocilia tips similarly to **b** (open arrowheads). MYO10-MD distributes diffusely without accumulating at stereocilia tips before AP21987 treatment (arrowhead) but accumulates at stereocilia tips with FRB-PST-EGFP after AP21987 treatment (arrows and open arrows). The amounts of MYO10-MD and PST at stereocilia tips are increased compared with untreated cells suggesting that MYO10-MD moved using PST as a scaffold. Bars, 5 µm. **d**, Single-molecule microscopy of MYO7A-HMM tethered to F-actin. Staircase-like trajectories consistent with step-wise movements are observed in the kymogram (arrows). The molecule indicated by arrows is shown in time-lapse images (magenta circles). AP21987, 500 nM. Single-plane time-lapse, every 1 s. Bars, 5 µm (cell image); 20 s and 2 µm (kymogram). **e,** Single-molecule microscopy of MYO10-MD tethered to F-actin. MYO10-MD also shows directional movements similarly to MYO7A-HMM (arrows and open arrows in the kymogram). The molecule indicated by arrows is shown in time-lapse images (magenta circles). AP21987, 500 nM. Single-plane time-lapse, every 100 ms. Bars, 5 µm (cell image); 2 s and 2 µm (kymogram).

Single-molecule microscopy demonstrated that MYO7A-HMM can move in a stereocilium using F-actin as a scaffold, but the trajectories in kymograms are different from those of dimers (Fig. 2d and Fig. 5d). After AP21987 treatment, only a small number of MYO7A-HMM molecules moved directionally toward stereocilia tips (Fig. 5d, Movies S10). Movements of MYO7A-HMM molecules tethered to F-actin were step-wise as observed for those anchored to the plasma membrane. MYO10-MD molecules also showed step-wise movements after the AP21987 treatment (Fig. 5e, Movie S11). These results suggest that myosin molecules tethered to F-actin move only when the tail is released from F-actin. These observations are not consistent with an “inchworm-like” movement proposed by others^40, 69^ because the step-sizes were 100–200 nm, which is much larger than the size of heavy meromyosin (∼15 nm)^70^. The restricted movements of myosins tethered to F-actin may be advantageous to anchor the components of the tip-link complex, including MYO7A, in the UTLD after being transported on the F-actin core by MYO7A dimers.

## 3. Discussion

The methodology proposed here expands the targets of single-molecule microscopy in live cells to three-dimensional organelles beyond stereocilia to include cilia, microvilli, filopodia and their associated diseases^71–75^. In this study, we successfully visualized and localized single HaloTag-fused protein molecules in live hair cells and used our method to elucidate how a single unconventional myosin can traffic in a stereocilium using their motor activities. Stereocilia are intricate mechanosensors consisting of more than 500 different proteins including actin monomers, F-actin bundling proteins, a multitude of MET channel and tip-link components and unconventional myosins^76^. Among these molecules, unconventional myosins are crucial molecular players during the development and maintenance of functional stereocilia because they transport and anchor specific stereocilia components using their highly diverse tail domains. Some unconventional myosins also interact with phosphatidylinositol 4,5-bisphosphate (PIP2) in the plasma membrane and are involved in adaptation of the MET channels (MYO1C)^77^ and maintenance of stereocilia architecture (MYO6)^78^. While decades of studies have identified various “cargo” of unconventional myosins in stereocilia, less is understood about how each myosin traffics in a stereocilium including whether or not each myosin is dimerized (or oligomerized). Elucidating the mechanisms underlying trafficking of unconventional myosins will be a basis for approaching questions more closely related to clinical practice, such as (1) why variants of some myosins (MYO6 and MYO7A) are associated with both autosomal dominant or recessive nonsyndromic hearing loss^79–82^ including the possibility of dominant-negative effects and (2) how myosin function could be restored therapeutically, especially for variants causing autosomal dominant progressive loss of hearing. Our single-molecule microscopy is a methodology of choice to explore these open questions through *in vivo* real-time functional analyses.

We speculate that the phenotype of constitutively active MYO7A mutants can be a good starting point to elucidate the formation of tip-links and the UTLD in developing stereocilia. MYO7A localizes in the UTLD region and tethers tip-links consisting of PCDH15 dimers and CDH23 dimers to the F-actin core on the CDH23 side^43, 83, 84^ (Fig. 6a). Two scaffolding proteins, Harmonin and SANS, bridge the interaction between MYO7A and CDH23^62–64^ and help to form a network of interactions (Fig. 6b). Harmonin is expressed in several isoforms in hair cells, which are collectively referred to as Harmonin b harboring the PST domain to interact with F-actin and as Harmonin a and c lacking the PST domain^42, 85^. Processive movements of MYO7A-RK/AA and MYO7A-ΔSH3-ΔM/F2 (Fig. 3) suggest that MYO7A can dimerize (or perhaps oligomerize) when the tail-mediated autoinhibition^47, 48^ is disabled. Cargo binding is one major mechanism to unleash MYO7A from the autoinhibitory state^53^ although further analyses are required to elucidate how MYO7A can be dimerized in stereocilia. SANS may be involved in this activation through the interaction with the first MyTH4-FERM domain (M/F1) because MYO7A-HMM-FKBP did not show movements without AP20187 at a detectable frequency (Fig. 2f). A previous study indicates that Harmonin is transported by MYO7A and SANS since localization of Harmonin b at stereocilia tips is lost in mice with defective MYO7A (*Myo7a^4626SB/4626SB^*) or SANS (*Ush1g^js/js^*)^45^. In addition, MYO7A-HMM did not move efficiently when anchored to the plasma membrane or tethered to F-actin (Figs. 4 and 5), which is consistent with previous studies showing that intact tip-links are not required to localize MYO7A and Harmonin b at stereocilia tips^45, 86^ and that the PST domain of Harmonin b is not necessary to form tip-links^42^. However, several studies suggest that there are alternative pathways to localize components of tip-links and the UTLD. Localization of CDH23 and PCDH15 in stereocilia tips, as well as that of MYO7A, are not lost in mice lacking Harmonin isoforms (*Ush1c*^−/−^)^45^. SANS can still localize in stereocilia of *Myo7a^4626SB/4626SB^* and in *Ush1c*^−/−^ mice although its localization is not limited to the UTLD^87^ and different from that in wild-type mice.

**Fig. 6:**
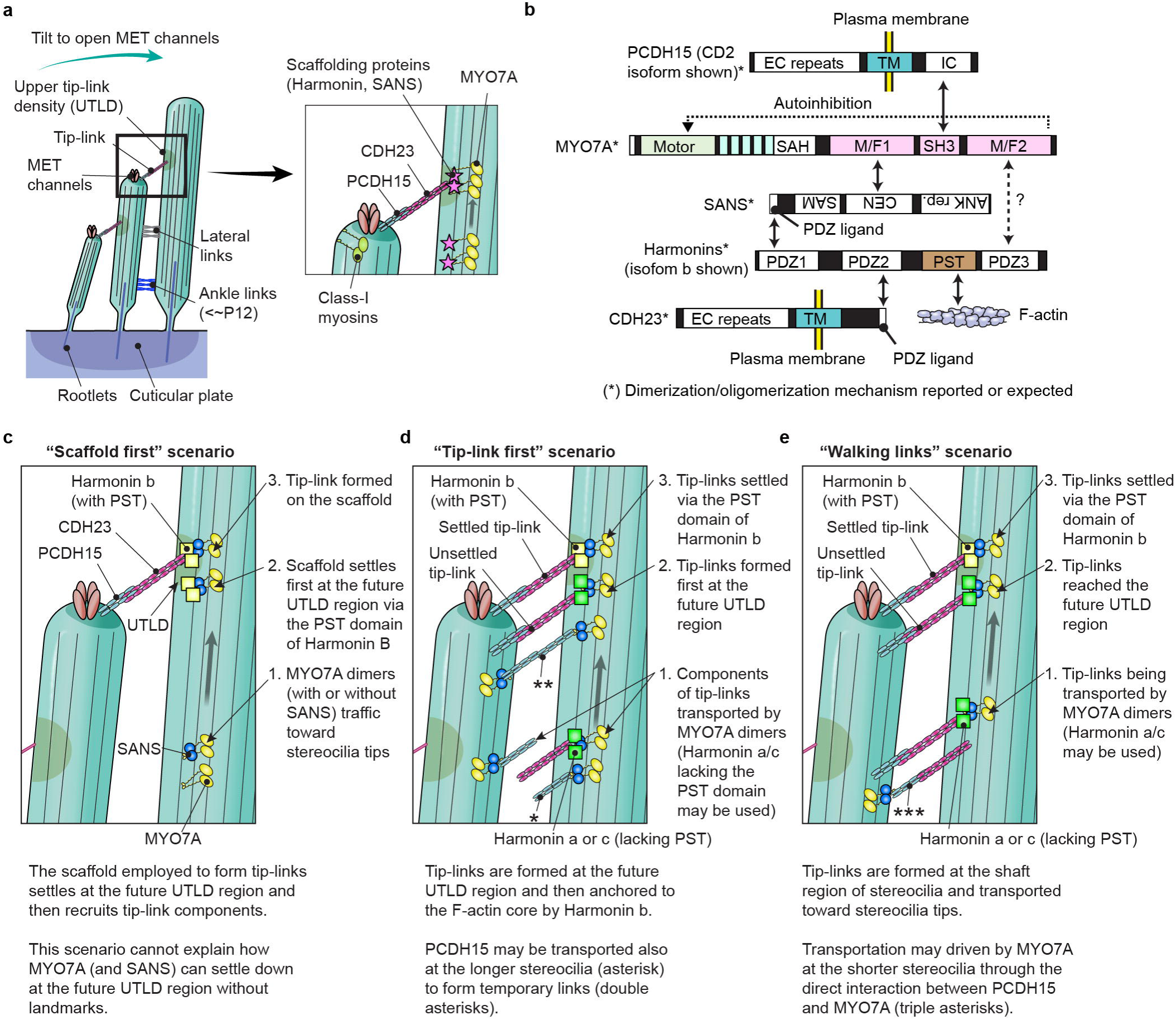
Possible scenarios for MYO7A-driven cargo transport in stereocilia. **a,** Architecture of stereocilia and the MET and tip-link complexes. The MET channels are localized at distal ends of stereocilia and physically gated by extracellular tip-links connected to the sides of adjacent stereocilia in the longer row at the region referred to as the upper tip-link density (UTLD). Each stereocilium contains a core of tightly-packed unidirectional F-actin whose diameter narrows near the hair cell apical surface (the taper region) and connects to the cuticular plate actin meshwork by a rootlet of more tightly packed F-actin. MYO7A is localized at the UTLD and involved in localization of components of the tip-link and the MET channel. Some class-I myosins are reported to play a role in adaptation during sustained sound stimulation. **b,** Major interacting partners of MYO7A in stereocilia. SANS and Harmonin bridge interactions with other partners. SANS is drawn upside down because it binds to the first MyTH4-FERM domain of MYO7A in the opposite direction^112^. The PST domain of Harmonin b can bind to F-actin^42^. The SAH domain of MYO7A has weak dimerization activity^53, 113^. SANS, PCDH15 and CDH23 can dimerize with each other ^64, 83, 84^ and may keep multiple MYO7A molecules in proximity. Oligomerization has been reported for Harmonin^63^. PCDH15 and CDH23 each have one transmembrane motif (TM)^83, 84^. The SH3 domain of MYO7A can interact with the intracellular portion (IC) of PCDH15 (CD2 isoform shown)^65^. **c–e**, Possible scenarios of MYO7A-driven localization and formation of tip-links and the UTLD. The “scaffold first” scenario (**c**) assumes that MYO7A (and SANS) traffics toward stereocilia tips, settles at the future UTLD region via the PST domain of Harmonin b and then recruits tip-link components. However, this scenario cannot explain how MYO7A (and SANS) can settle down at the future UTLD region without landmarks. The “tip-link first” scenario (**d**) assumes that tip-link components are transported toward stereocilia tips and then form tip-links at the UTLD region. Different isoforms of Harmonin might be used during the transport (Harmonin a or c lacking the PST domain) and after the transport (Harmonin b with the PST domain) because MYO7A anchored to the F-actin by the PST domain of Harmonin b cannot traffic efficiently (see Fig. 5). PCDH15 may be transported in both shorter and longer stereocilia and form temporary links as indicated by BAPTA-mediated remodeling experiments^88^. The “walking links” scenario (**e**) supported by a previous cryo-electron microscopy study assumes that tip-links are formed on the shaft of stereocilia and then transported toward stereocilia tips^89^. The “walking links” and “tip-link first” scenarios may coincide because many uncoupled PCDH15 molecules were reported at stereocilia tips in the same study^89^.

Our data support several scenarios that might localize tip-link components: (1) scaffolds of MYO7A, SANS and Harmonin b settle on the F-actin core and then recruit tip-link components (Fig. 6c, “scaffold first” scenario), (2) tip-links are formed at the future UTLD region and then are anchored to the F-actin core by Harmonin b (Fig. 6d, “tip-link first” scenario), and (3) fully formed tip-links are transported toward stereocilia tips and are then anchored to the F-actin core by Harmonin b (Fig. 6e, “walking links” scenario). In these scenarios, SANS, PCDH15 and CDH23 may dimerize MYO7A since these proteins can homodimerize with each other^64, 83, 84^. Harmonin b might also be another mediator in dimerization or oligomerization through its oligomerization activity^63^. Among these three scenarios, the “scaffold first” scenario is less likely because it cannot explain how MYO7A (and SANS) can settle down at the future UTLD region without a landmark. The “tip-link first” scenario can be supported by remodeling experiments of tip-links because tip-links are newly formed after being disassembled by BAPTA-mediated extracellular calcium chelation^88^. This study also reports that temporary links consisting of only PCDH15 are formed after BAPTA treatment (Fig. 6d, double asterisks) subsequently replacing PCDH15 on the taller stereocilia side with CDH23 to form mature tip-links^88^. In contrast, the “walking links” scenario is supported by a previous cryoelectron microscopy study using anti-PCDH15 antibodies, which detected multiple “lateral links” at the shaft region of stereocilia^89^. These links consist of PCDH15 (∼50 nm long) on one side and a longer partner on the other side (∼120 nm long), putatively CDH23^89^. However, the “tip-link first” and “walking links” scenarios may coincide because many uncoupled PCDH15 molecules are reported to localize near stereocilia tips in the cryoelectron microscopy study^89^. These uncoupled PCDH15 molecules may be actively transported, for example, by the interaction with the SH3 domain of MYO7A^65^ (Fig. 6d, asterisk) and also provokes the idea that active transport of fully-formed tip-links may occur on the PCDH15 side (Fig. 6e, triple asterisks). In addition, our study shows that movements of MYO7A is restricted when its tail is anchored to the F-actin core by the PST domain of Harmonin b (Fig. 5). Harmonin a and c lacking the PST domain should be advantageous for effective active transport on the CDH23 side (Fig. 6, d and e). Further analyses, especially on trafficking of PCDH15 and CDH23, are necessary to conclude which circumstance most likely explains how tip links arise *in vivo*.

In addition to MYO7A, our single molecule imaging studies can be applied to other unconventional myosins functioning in F-actin protrusions to answer unresolved questions. For example, it is still uncertain how class-III myosins move in F-actin protrusions including stereocilia. Currently, there is no evidence that MYO3A has a dimerization sequence, such as a coiled-coil domain^90^. Instead, MYO3A can move directionally in F-actin protrusions using the THDI and THDII domains in its tail, which interact with F-actin binding proteins, ESPN isoform 1 or ESPNL, and F-actin, respectively^39, 40, 91^. Our method may elucidate how MYO3A can move toward the barbed ends when their tail is tethered to F-actin and how directional movements of MYO3A is regulated by the autoinhibitory kinase domain^92^. Stepwise movements of MYO7A-HMM and MYO10-MD tethered to F-actin suggest that MYO3A may traffic in stereocilia similarly, not by “inchworm-like” movements as previously proposed^40, 69^. In addition, MYO10-MD can move in stereocilia using various scaffolds and even as a monomer (Figs. 4e and 5e). This observation gives us a clue as to why MYO7A and MYO7B are utilized in stereocilia and microvilli, both of which are more stable and long-lived than filopodia^93^. It is known that recruitment of MYO10 to the plasma membrane can induce formation of filopodia-like F-actin protrusions^66^. Movements of MYO7A are restricted when anchored to plasma membranes, which might prevent MYO7A from forming unwanted F-actin protrusions that would destroy the stereocilia architecture. MYO10-MD tethered to the plasma membrane or F-actin accumulates at stereocilia tips and can alter the architecture of stereocilia (Figs. 4c and 5c).

Our methodology enables functional analyses of protein molecules in the context of live cells. Protein kinetics in stereocilia are complex due to the multitude of components and to the limited diffusion from the tightly-packed F-actin and the enveloping plasma membrane^49^. Moreover, the behavior of proteins *in vivo* can be completely different from that *in vitro* as we demonstrated for CapZ, an F-actin barbed end capper, in lamellipodia^94^. In this study, we noticed that MYO7A-HMM dimers move 10-fold faster than in a previous study^58^. Although this difference can be partially attributed to the difference in temperature (37°C in this study vs. 25°C in the previous study^58^), a previous MYO10 dimer study shows only less than 2-fold increase in velocity between 25°C and 37°C^59^. In filopodia, it was shown that unidirectionally bundled F-actin is advantageous for MYO7A-HMM dimers to move rapidly without changing the “actin tracks”, but the increase in velocity is less than 2-fold^58^. One possibility is that some dimers “jump” toward the barbed ends between two processive movements or binding events using restricted diffusion in stereocilia^49^. Further analyses are required to test this hypothesis. In addition, analyses of molecular turnover in a semi-equilibrium state could also benefit from our technique. Actin and regulatory proteins are replenished in the F-actin core while the shape, width and length of stereocilia remain largely unchanged on the apical surface of hair cells^12, 24, 95^. Our methodology may answer another crucial question as to how the architecture of stereocilia remains stable for the lifespan of a normal hearing person^96^.

In this study, we used a symmetrical diSPIM microscope equipped with two 40×, 0.8 NA water-immersion objective lenses^26^ used for single dye detection in our previous study^12^. Our method could be improved by imaging with a higher numerical aperture (NA) objective to increase localization precision and light collection. Improved NA could enable richer analyses of myosin trafficking in stereocilia including investigation of their step-sizes^97^ and processivity and also may allow single-molecule microscopy in stereocilia of cochlear hair cells. Step-size analysis could be facilitated in future studies by using an asymmetric diSPIM design, featuring higher NA lenses and a suitably long working distance^98^. Another future improvement would be to avoid the necessity for an activation mechanism. This study employed chemically-induced dimerization techniques to “turn on” myosin trafficking after cells were ready for imaging. Methods for direct delivery of fluorescently-labeled protein molecules, such as microinjection, will be useful for this purpose and also for visualizing replenishment and dissociation of protein molecules in stereocilia avoiding the complexities associated with blinking fluorescent dyes^99,100^.

Single-molecule microscopy enables real-time observation of molecules at work, such as unconventional myosins trafficking in stereocilia, as we demonstrate in this study. Recent advances in light-sheet microscopy allowed us to apply this powerful technique to stereocilia, which had been challenging to image due to their architecture protruding upward from the apical cell surface. Although mechanosensory stereocilia are unique to the inner ear, our data should be applicable to analyze active cargo transport in other F-actin protrusions, microvilli and filopodia, where diffusion of proteins are restricted^49^. In addition, the flexible spatial arrangement in our methodology will be useful for other three-dimensional organs and organelles including primary cilia, kinocilia, migrating cells and perhaps neuronal cell layers. We speculate that our methodology will contribute to expanding the target of single-molecule fluorescence microscopy and help formulating therapeutic strategies not only for sensorineural hearing loss but also for other diseases, such as ciliopathies^71, 72^, neurodegenerative diseases^101^, inflammatory bowel diseases^73, 74^ and cancer^75^.

## 4. Online Methods

### 4.1. Plasmids and cDNAs

To express HaloTag-fused proteins, we modified the pEGFP-C1 (Clonetech) vector by replacing the EGFP sequence with a HaloTag sequence that was PCR amplified from the pHTC HaloTag® CMV-neo (Promega). Two silent mutations were introduced to the wild-type HaloTag sequence to disable the XhoI and SalI endonuclease restriction sites. The resulting vector pHaloTagXS-C1 was used to express HaloTag-fused proteins. A plasmid encoding HaloTag-fused human β-actin was constructed using this vector and the actin sequence deriving from pEGFP-actin (Clonetech). Plasmids encoding HaloTag-fused MYO7A fragments were constructed using the pHaloTagXS-C1 vector and inserts amplified from a plasmid encoding EGFP-fused mouse *Myo7a* isoform 1 (NM_001256083.1)^36^, a gift from Erich Boger, NIDCD. Plasmids encoding HaloTag-fused MYO10 fragments were constructed similarly using pcDNA3.1 Zeo+ EGFP-Myo10noCC-FRB-myc^66^, a gift from Matthew Tyska, Vanderbilt University. To express HaloTag-fused myosin fragments with the C-terminal FKBP (e.g., HaloTag-MYO7A-HMM-FKBP), a DNA fragment encoding FKBP (LC087168) was amplified from GFP-M7HMM-FKBP^53^, a gift from Mitsuo Ikebe, University of Texas Tyler Health Center, with the C-terminal HA tag originally in GFP-M7HMM-FKBP. Expression of HaloTag-fused myosin fragments with a C-terminal FRB (e.g., HaloTag-MYO7A-HMM-FRB) was achieved by replacing the FKBP portion of HaloTag-fused myosin fragments with the C-terminal FKBP (e.g., HaloTag-MYO7A-HMM-FKBP) with the FRB sequence of pEGFP-FRB^102^ (Addgene #25919). Thus, the C-terminal FKBP and FRB attached to HaloTag-fused myosin fragments have an additional HA tag in this study. A plasmid to express IL2Rα-EGFP-FKBP was constructed from pEGFP-C1 inserting a DNA fragment encoding the Tac antigen (human IL2 receptor alpha subunit; NM_000417.3) amplified from TAC-GFP^103^ (Addgene # 162494) and the FKBP fragment. A mouse USH1C fragment (residues 296–728 of NM_01163733) was expressed with FRB at the N-terminus and EGFP at the C-terminus using a plasmid constructed from pEGFP-N3 (Clonetech) inserting DNA fragment amplified from pEGFP-FRB^102^ and a USH1C fragment amplified from the cDNA of USH1C isoform b4, a gift from Nicolas Grillet at Stanford University.

### 4.2. Animals

All animal experiments were performed in accordance with the National Institutes of Health Guidelines for the Care and Use of Laboratory Animals and approved by the Animal Care and Use Committees at the NIH (No. 1263 to TBF). Mouse neonates were obtained from timed pregnant C57BL6/J females purchased from the Jackson Laboratory or from our in-house C57BL6/J colony. Vestibular sensory epithelia were harvested from neonates at postnatal day (P) 2–5 after being euthanized by decapitation.

### 4.3. Explant culture and transfection of vestibular sensory epithelia

Inner ear sensory epithelia were cultured and transfected using a Helios® Gene Gun System (Bio-Rad) as previously described with slight modification^51^. Briefly, utricles and saccules of P2– 5 mice were isolated in Leibovitz’s L-15 medium (Thermo Fisher) after removing the otoliths using a 30G needle. Isolated sensory epithelia were placed on a glass-bottom dish (MatTek Corporation) coated with rat-tail collagen I (A1048301, Thermo Fisher) matrix and cultured in Dulbecco’s Modified Eagle Medium/Nutrient Mixture F-12 (DMEM/F12, Thermo Fisher) supplemented with 7% FCS (Atlanta Biologicals) and 20 µg/mL ampicillin (Sigma) at 37°C in 5% CO_2_. After culturing for 6–20 h, vestibular hair cells were transfected using a gene-gun and 1.0 µm Gold Microcarriers (1652263, Bio-Rad) propelled by helium pulses at 110–115 psi. Gold microcarriers were coated with a 1:3 or 1:1 mixture of two plasmids encoding a HaloTag-fused protein and EGFP (or an EGFP-fused protein), respectively.

### 4.4. Single molecule microscopy of live hair cell stereocilia

Vestibular sensory epithelia co-expressing a HaloTag-fused protein and EGFP (or EGFP-fused protein) were labeled using 0.01–1 nM JFX554-conjugated HaloTag ligands^52^ (gift from Luke Lavis at Janelia Research Campus) diluted in culture medium (DMEM/F12 supplemented with 7% FCS and 20 µg/mL Ampicillin) for 30 min at 37°C in 5% CO_2_. Unreacted HaloTag ligands were removed by washing the tissue in the culture medium for 3–5 s three times and by incubating samples at 37°C in 5% CO_2_ up to 6 h until live imaging was performed. Samples on a collagen matrix were detached from glass-bottom dishes with the underneath coverslip, mounted in a 10-cm plastic culture dish on a 1–2 mm droplet of vacuum grease (Fisher Scientific) and then incubated in Leibovitz’s L-15 Medium without phenol red (21083027, Thermo Fisher) warmed at 37°C. For conditional dimerization, the medium was gently removed using plastic transfer pipettes and replaced with a new L-15 medium containing 200 nM AP20187 (Sigma) or 500 nM AP21987 (Takara Bio).

Images were acquired using a custom-made symmetrical dual-view inverted selective plane illumination microscope (diSPIM)^26^ installed in a 37°C incubator (Applied Scientific Instrumentation). The diSPIM was equipped with 40× Nikon CFI APO NIR objectives (0.80 NA, 3.5 mm WD; Nikon), ORCA-Fusion Digital CMOS cameras (C14440-20UP; Hamamatsu), an OBIS 488 nm LX 150 mW Laser (Coherent), an OBIS 561 nm LS 150 mW Laser (Coherent) and a W-VIEW GEMINI Image splitting optic (Hamamatsu) with a 561 nm laser BrightLine single-edge super-resolution/TIRF dichroic mirror (Semrock), a 525/50 nm BrightLine single-band bandpass filter (Semrock) and a 568 nm EdgeBasic best-value long-pass edge filter (Semrock). The microscope was controlled by Micro-Manager (https://micro-manager.org/), a plugin for ImageJ^104–106^.

The entire architecture of stereocilia was visualized using fluorescence from co-expressed EGFP or EGFP-fused proteins by a volume scan of 0.5-µm thickness illuminating the 488-nm laser at approximately 0.01 kW/cm^2^ for 100 ms per slice. HaloTag-fused protein molecules labeled with JFX554 were visualized illuminating the 561-nm laser at approximately 0.2 kW/cm^2^ for 100 ms per plane. HaloTag-actin in Fig. 1, **a** and **b**, was imaged by a single volume scan of 0.5-µm thickness. Other images were acquired every 0.1 to 1 s by single-plane time-lapse imaging. Sample drift was corrected using our custom-made Python scripts available at GitHub (http://github.com/takushim/momomagick) implementing phase-only correlation with subpixel matching^107^ and least-square image matching^108^. Fluorescent puncta were manually tracked using our custom-made Python script with a graphical user interface (http://github.com/takushim/momotrack). The point spread function of the 40× objective lens was obtained from a previous study^109^. Kymograms were generated using the Fiji platform^110^.

### 4.5. Fluorescence histochemistry

Explant cultures of vestibular sensory epithelia were fixed in 4% paraformaldehyde (PFA; Electron Microscopy Sciences) in PBS for 30 min at room temperature (RT) and washed in PBS. For conditional dimerization, samples were treated with 100 nM AP20187 (Sigma) or 500 nM AP21987 (Takara Bio) for 2 h. The concentration of AP20187 was lowered because 200 nM AP21087 caused strong accumulation of MYO7A-HMM dimers at stereocilia tips and often damaged the architecture of stereocilia. Samples were permeabilized and blocked in PBS containing 1% bovine serum albumin (BSA) and 0.2% Triton-X100 for 20–30 min at RT. HaloTag-fused proteins were visualized by reacting with 200 nM JFX554-HaloTag-ligand^52^ (gift from Luke Lavis) in PBS with 0.2% Triton-X 100 for 1 h at RT. F-actin was visualized by supplementing the HaloTag ligand solution with 10–30 nM of Alexa Fluor™ Plus 405 Phalloidin (Thermo Fisher). Confocal images were acquired using a Zeiss LSM880 with Airyscan processing (Zeiss).

## 5. Acknowledgements

We thank Drs. Dennis Winkler and Mhamed Grati for valuable comments; Elizabeth Bernhard, Sherly Michel and Alexander Callahan for managing mouse colonies; Jiji Chen and Min Guo for comments on single-molecule microscopy and image processing; and Erina He for her beautiful diagrams. Images using the diSPIM were acquired in the Advanced Imaging and Microscopy Resource of the NIBIB. This article is subject to Howard Hughes Medical Institute (HHMI)’s Open Access to Publications policy. HHMI laboratory heads have previously granted a non-exclusive CC BY 4.0 license to the public and a sub-licensable license to HHMI in their research articles. Pursuant to those licenses, the author-accepted manuscript of this article can be made freely available under a CC BY 4.0 license immediately upon publication.

## 6. Author Contributions

**Takushi Miyoshi:** Conceptualization, Methodology, Software, Validation, Investigation, Writing – Original draft, Visualization, Project administration and Funding acquisition. **Harshad D. Vishwasrao:** Methodology and Investigation. **Inna A. Belyantseva:** Methodology and Writing – Review & Editing and Visualization. **Mrudhula Sajeevadathan:** Validation, Writing – Review & Editing and Visualization. **Yasuko Ishibashi:** Validation and Writing – Review & Editing. **Samuel Adadey:** Validation and Writing – Review & Editing. **Narinobu Harada:**

Conceptualization and Methodology. **Hari Shroff:** Conceptualization, Methodology, Writing – Review & Editing, Supervision and Funding acquisition. **Thomas B. Friedman:** Writing – Review & Editing, Supervision, Project administration and Funding acquisition.

## Supplemental Figures

**Fig. S1:**
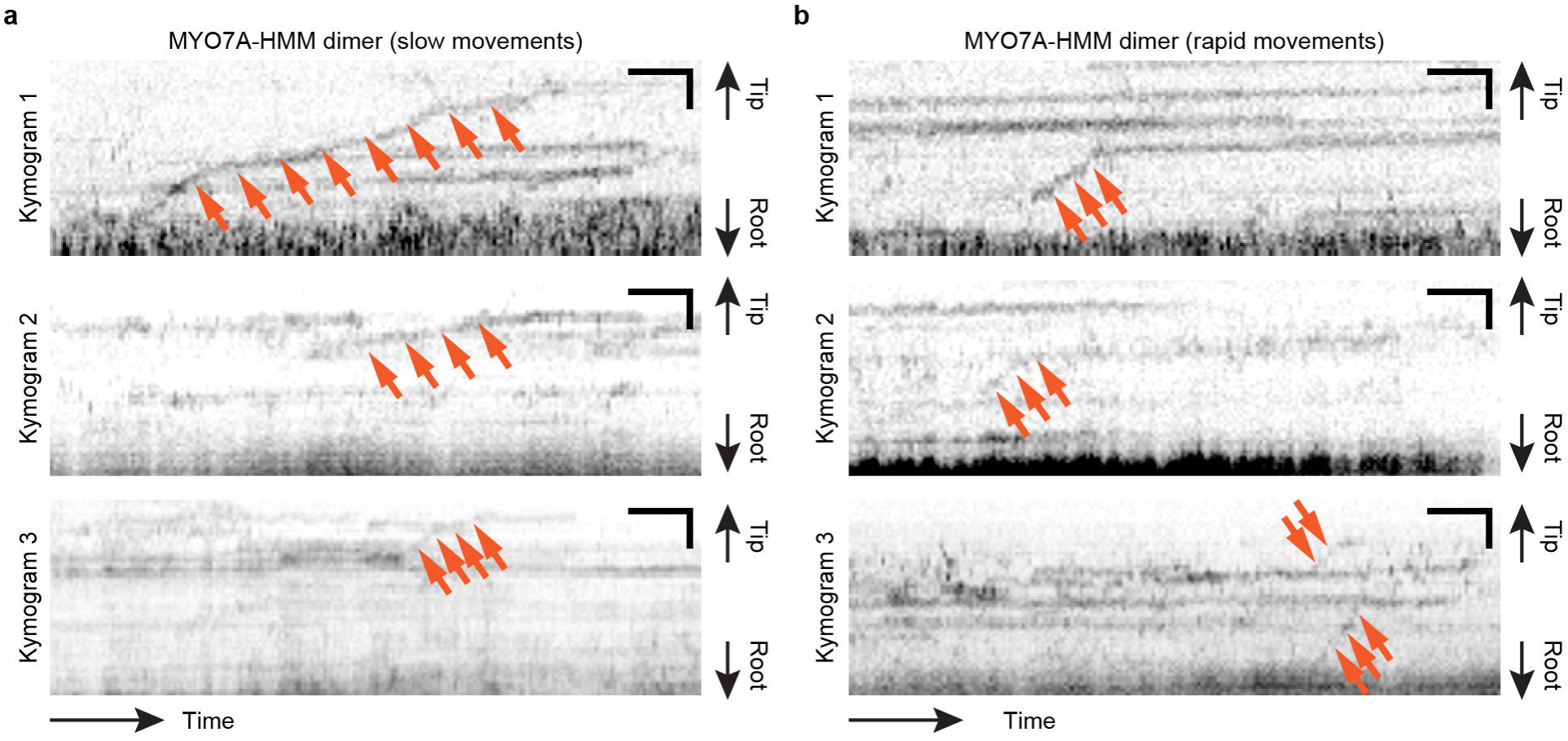
Processive movements of MYO7A-HMM dimers in stereocilia. Representative kymograms of HaloTag-MYO7A-HMM-FKBP in cells treated with 200 nM AP20187. Trajectories of slow movements (**a**) and rapid movements (**b**) are shown (arrows). Single-plane time-lapse, every 1 s. Bars, 2 µm and 20 s.

**Fig. S2:**
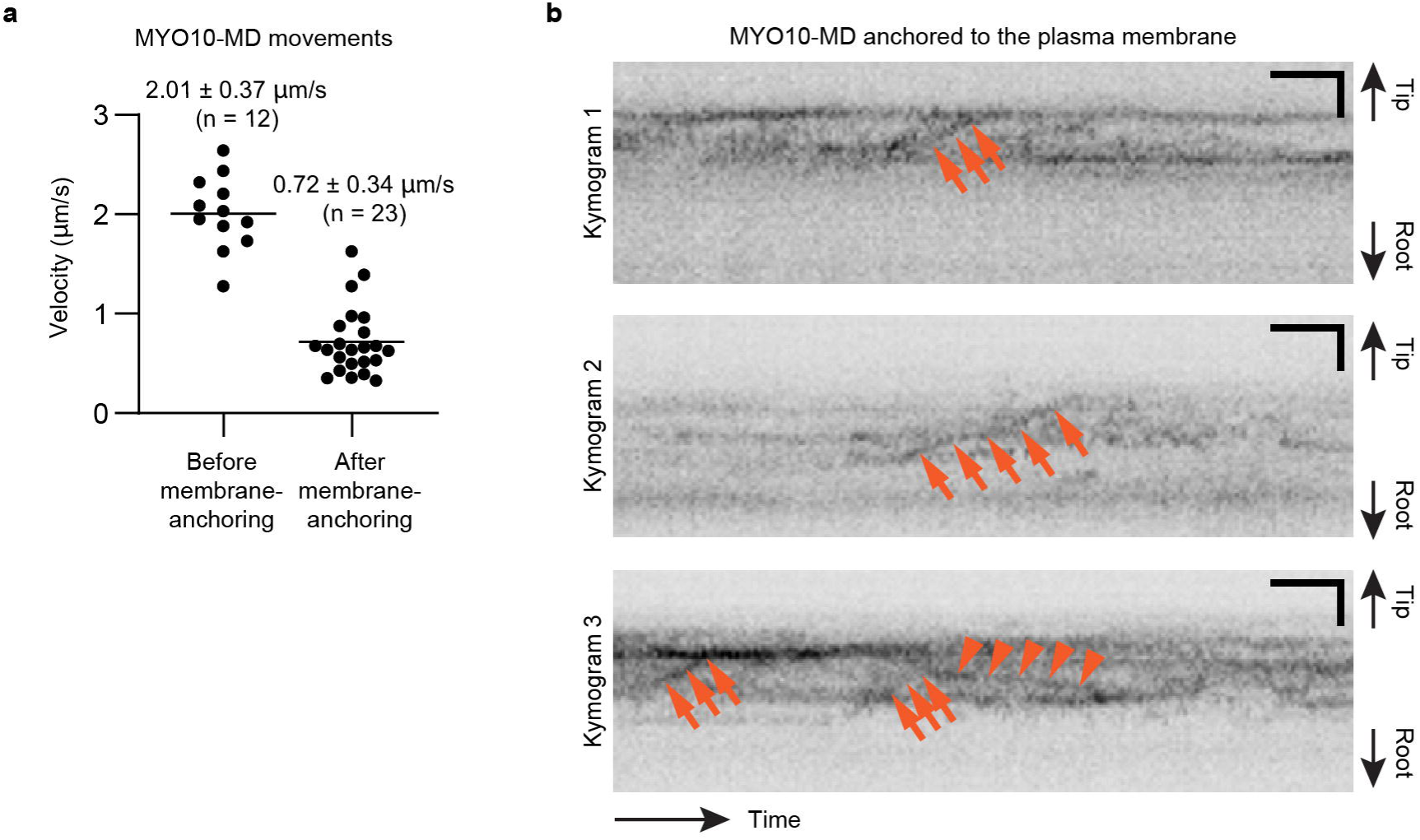
Movements MYO10-MD anchored to the plasma membrane of stereocilia. **a**, Velocities of MYO10 molecules moving in stereocilia before and after the 500 nM AP21987 treatment. The mean velocity was 2.01 ± 0.37 µm/s (n = 12, mean ± standard deviation) before dimerization and 0.72 ± 0.34 µm/s (n = 23) after dimerization. **b**, Representative kymograms of membrane-anchored MYO10-MD molecules. Continuous trajectories are consistent with processive and directional movements (arrows). Retrograde movements are also observed (arrowheads). Single-plane time-lapse, every 100 ms. Bars, 2 µm and 6 s.

## Supplemental information

**Movie S1: Time-lapse images of HaloTag-actin**

Vestibular hair cell (P2) expressing HaloTag-actin was imaged as a control for proteins stably bound to the F-actin core. Most of the fluorescent puncta remain in the same location and disappear suddenly due to photobleaching or transition to the dark state (representatively indicated by magenta circles). JFX554, 0.01 nM. Single-plane time-lapse, every 1 s. Exposure, 100 ms at 0.2 kW/cm^2^. Bar, 5 µm.

**Movie S2: Time-lapse images of non-fused HaloTag**

Vestibular hair cell (P2) expressing non-fused HaloTag was imaged as a control for diffusing proteins. Most fluorescent puncta disappear after one frame. JFX554, 0.1 nM. Single-plane time-lapse, every 1 s. Exposure, 100 ms at 0.2 kW/cm^2^. Bar, 5 µm.

**Movie S3: Time-lapse images of HaloTag-MYO7A-HMM-FKBP with the AP20187 treatment**

Vestibular hair cell (P2) expressing HaloTag-MYO7A-HMM-FKBP was imaged after adding 200 nM AP20187 to the culture medium. Molecules showing directional movements are indicated by magenta circles. JFX554, 0.3 nM. Single-plane time-lapse, every 1 s. Exposure, 100 ms at 0.2 kW/cm^2^. Bar, 5 µm.

**Movie S4: Time-lapse images of HaloTag-MYO7A-HMM-FKBP without AP20187 treatment**

Vestibular hair cell (P2) expressing HaloTag-MYO7A-HMM-FKBP was imaged without adding AP21087 to the culture medium. No directional movements are observed. JFX554, 0.3 nM. Single-plane time-lapse, every 1 s. Exposure, 100 ms at 0.2 kW/cm^2^. Bar, 5 µm.

**Movie S5: Time-lapse images of HaloTag-MYO7A-RK/AA**

Vestibular hair cell (P2) expressing HaloTag-MYO7A-RK/AA, which has two missense mutations (p.R2127A and p.K2130A) disabling autoinhibition of the motor domain, was imaged every 1 s by single-plane time-lapse acquisition. Molecules showing directional movements are indicated by magenta circles. JFX554, 0.3 nM. Exposure, 100 ms at 0.2 kW/cm^2^. Bar, 5 µm.

**Movie S6: Time-lapse images of HaloTag-MYO7A-ΔSH3-ΔM/F2**

Vestibular hair cell (P2) expressing HaloTag-MYO7A-ΔSH3-ΔM/F2, whose tail is truncated to disable autoinhibition of the motor domain, was imaged every 1 s by single-plane time-lapse acquisition. A molecule showing directional movements is indicated by magenta circles. JFX554, 0.3 nM. Exposure, 100 ms at 0.2 kW/cm^2^. Bar, 5 µm.

**Movie S7: Time-lapse images of membrane-anchored HaloTag-MYO7A-HMM-FRB**

Vestibular hair cell (P2) co-expressing HaloTag-MYO7A-HMM-FRB and IL2Rα-EGFP-FKBP. The cell is treated with 500 nM AP21987 to anchor MYO7A-HMM to the plasma membrane. Molecules showing stepwise, directional movements toward stereocilia tips are indicated by magenta circles. Single-plane time-lapse, every 300 ms. JFX554, 0.3 nM. Exposure, 100 ms at 0.2 kW/cm^2^. Bar, 5 µm.

**Movie S8: Time-lapse images of HaloTag-MYO10-MD-FRB before membrane anchoring**

Vestibular hair cell (P2) co-expressing HaloTag-MYO10-MD-FRB and IL2Rα-EGFP-FKBP. The cell is imaged without the AP21987 treatment. A small number of molecules show rapid directional movements toward stereocilia tips (magenta circles). Single-plane images are acquired every 100 ms. JFX554, 0.3 nM. Exposure, 100 ms at 0.2 kW/cm^2^. Bar, 5 µm.

**Movie S9: Time-lapse images of HaloTag-MYO10-MD-FRB after membrane anchoring**

Vestibular hair cell (P2) co-expressing HaloTag-MYO10-MD-FRB and IL2Rα-EGFP-FKBP. The cell is treated with 500 nM AP21987 to anchor MYO10-MD to the plasma membrane. Molecules showing processive movements are indicated by magenta circles. Single-plane images are acquired every 100 ms. JFX554, 0.3 nM. Exposure, 100 ms at 0.2 kW/cm^2^. Bar, 5 µm.

**Movie S10: Time-lapse images of HaloTag-MYO7A-HMM-FKBP tethered to F-actin**

Vestibular hair cell (P2) co-expressing HaloTag-MYO7A-HMM-FKBP and FRB-PST-EGFP. The cell is treated with 500 nM AP21987 to bind the C-terminus of MYO7A-HMM to F-actin. A small number of molecules show stepwise, directional movements toward stereocilia tips (magenta circles). Single-plane time-lapse, every 1 s. JFX554, 0.3 nM. Exposure, 100 ms at 0.2 kW/cm^2^. Bar, 5 µm.

**Movie S11: Time-lapse images of HaloTag-MYO10-MF-FKBP tethered to F-actin**

Vestibular hair cell (P2) co-expressing HaloTag-MYO10-MD-FKBP and FRB-PST-EGFP. The cell is treated with 500 nM AP21987 to bind the C-terminus of MYO10-MD to F-actin. Molecules start stepwise, directional movements toward stereocilia tips (magenta circles). Single-plane time-lapse, every 100 ms. JFX554, 0.3 nM. Exposure, 100 ms at 0.2 kW/cm^2^. Bar, 5 µm.

## Abbreviations

GFP: green fluorescent protein
EGFP: enhanced green fluorescent protein
F-actin: filamentous actin
diSPIM: dual-view inverted selective plane illumination microscope
DFNA: locus for dominantly inherited nonsyndromic hearig loss
DFNB: locus for recessively inherited nonsyndromic hearing loss
MET: mechanoelectrical transduction
NA: numerical aperture
UTLD: upper tip-link density
ATP: adenosine triphosphate.

## References

1 Olusanya, B. O., Davis, A. C. & Hoffman, H. J. Hearing loss: rising prevalence and impact. Bull World Health Organ 97, 646–646A (2019). 10.2471/BLT.19.224683

2. Tanna, R. J., Lin, J. W. & De Jesus, O. Sensorineural Hearing Loss, <https://www.ncbi.nlm.nih.gov/pubmed/33351419> (2024).

3 Gillespie, P. G. & Muller, U. Mechanotransduction by hair cells: models, molecules, and mechanisms. Cell 139, 33–44 (2009). 10.1016/j.cell.2009.09.010

4 Eatock, R. A. & Lysakowski, A. in *Vertebrate Hair Cells Springer Handbook of Auditory Research* (eds Ruth Anne Eatock, Richard R. Fay, & Arthur N. Popper) Ch. Chapter 8, 348–442 (Springer New York, 2006).

5 Wong, A. C. & Ryan, A. F. Mechanisms of sensorineural cell damage, death and survival in the cochlea. Front Aging Neurosci 7, 58 (2015). 10.3389/fnagi.2015.00058

6 Lv, J. et al. AAV1-hOTOF gene therapy for autosomal recessive deafness 9: a single-arm trial. Lancet (2024). 10.1016/S0140-6736(23)02874-X

7 Qi, J. et al. AAV-Mediated Gene Therapy Restores Hearing in Patients with DFNB9 Deafness. Adv Sci (Weinh) 11, e2306788 (2024). 10.1002/advs.202306788

8 Wang, H. et al. Hair cell-specific Myo15 promoter-mediated gene therapy rescues hearing in DFNB9 mouse model. Mol Ther Nucleic Acids 35, 102135 (2024). 10.1016/j.omtn.2024.102135

9 Duan, M., Venail, F., Spencer, N. & Mezzina, M. Treatment of peripheral sensorineural hearing loss: gene therapy. Gene Ther 11 **Suppl 1**, S51–56 (2004). 10.1038/sj.gt.3302369

10 Heller, I., Dulin, D. & Peterman, E. J. G. *Single molecule analysis : methods and protocols*. Third edition edn, (Humana Press, 2024).

11 Liu, Z., Lavis, L. D. & Betzig, E. Imaging live-cell dynamics and structure at the single-molecule level. Mol Cell 58, 644–659 (2015). 10.1016/j.molcel.2015.02.033

12 Miyoshi, T. et al. Semi-automated single-molecule microscopy screening of fast-dissociating specific antibodies directly from hybridoma cultures. Cell Rep 34, 108708 (2021). 10.1016/j.celrep.2021.108708

13 Forster, T. Energiewanderung und fluoreszenz. Naturwissenschaften 33, 166–175 (1946).

14 Ashkin, A., Dziedzic, J. M., Bjorkholm, J. E. & Chu, S. Observation of a single-beam gradient force optical trap for dielectric particles. Opt Lett 11, 288–290 (1986). 10.1364/OL.11.000288

15 Magde, D., Elson, E. & Webb, W. W. Thermodynamic Fluctuations in a Reacting System---Measurement by Fluorescence Correlation Spectroscopy. Physical Review Letters 29, 705–708 (1972). 10.1103/PhysRevLett.29.705

16 Rust, M. J., Bates, M. & Zhuang, X. Sub-diffraction-limit imaging by stochastic optical reconstruction microscopy (STORM). Nature methods 3, 793–795 (2006). 10.1038/nmeth929

17 Heilemann, M. et al. Subdiffraction-resolution fluorescence imaging with conventional fluorescent probes. Angew Chem Int Ed Engl 47, 6172–6176 (2008). 10.1002/anie.200802376

18 Betzig, E. et al. Imaging intracellular fluorescent proteins at nanometer resolution. Science 313, 1642–1645 (2006). 10.1126/science.1127344

19 Jungmann, R. et al. Multiplexed 3D cellular super-resolution imaging with DNA-PAINT and Exchange-PAINT. Nature methods 11, 313–318 (2014). 10.1038/nmeth.2835

20 Schueder, F. et al. Universal Super-Resolution Multiplexing by DNA Exchange. Angew Chem Int Ed Engl 56, 4052–4055 (2017). 10.1002/anie.201611729

21 Elliott, A. D. Confocal Microscopy: Principles and Modern Practices. Curr Protoc Cytom 92, e68 (2020). 10.1002/cpcy.68

22 Li, X. et al. Three-dimensional structured illumination microscopy with enhanced axial resolution. Nat Biotechnol 41, 1307–1319 (2023). 10.1038/s41587-022-01651-1

23 Icha, J., Weber, M., Waters, J. C. & Norden, C. Phototoxicity in live fluorescence microscopy, and how to avoid it. Bioessays 39 (2017). 10.1002/bies.201700003

24 Drummond, M. C. et al. Live-cell imaging of actin dynamics reveals mechanisms of stereocilia length regulation in the inner ear. Nat Commun 6, 6873 (2015). 10.1038/ncomms7873

25 Dailey, M. E., Marrs, G. S. & Kurpius, D. Maintaining live cells and tissue slices in the imaging setup. Cold Spring Harbor protocols 2011, pdb top105 (2011). 10.1101/pdb.top105

26 Wu, Y. et al. Spatially isotropic four-dimensional imaging with dual-view plane illumination microscopy. Nat Biotechnol 31, 1032–1038 (2013). 10.1038/nbt.2713

27 McGrath, J. et al. Actin at stereocilia tips is regulated by mechanotransduction and ADF/cofilin. Curr Biol (2020). 10.1016/j.cub.2020.12.006

28 Weil, D. et al. Defective myosin VIIA gene responsible for Usher syndrome type 1B. Nature 374, 60–61 (1995). 10.1038/374060a0

29 Miyoshi, T., Belyantseva, I. A., Sajeevadathan, M. & Friedman, T. B. Pathophysiology of human hearing loss associated with variants in myosins. Front Physiol 15, 1374901 (2024). 10.3389/fphys.2024.1374901

30 Moreland, Z. G. & Bird, J. E. Myosin motors in sensory hair bundle assembly. Current opinion in cell biology 79, 102132 (2022). 10.1016/j.ceb.2022.102132

31 Sekerkova, G. et al. Espins are multifunctional actin cytoskeletal regulatory proteins in the microvilli of chemosensory and mechanosensory cells. The Journal of neuroscience : the official journal of the Society for Neuroscience 24, 5445–5456 (2004). 10.1523/JNEUROSCI.1279-04.2004

32 Corey, D. P. & Hudspeth, A. J. Ionic basis of the receptor potential in a vertebrate hair cell. Nature 281, 675–677 (1979). 10.1038/281675a0

33 Glowatzki, E. & Fuchs, P. A. Transmitter release at the hair cell ribbon synapse. Nat Neurosci 5, 147–154 (2002). 10.1038/nn796

34 Corey, D. P. & Hudspeth, A. J. Kinetics of the receptor current in bullfrog saccular hair cells. The Journal of neuroscience : the official journal of the Society for Neuroscience 3, 962–976 (1983). 10.1523/JNEUROSCI.03-05-00962.1983

35 Li, J. & Zhang, M. Cargo Binding by Unconventional Myosins. Adv Exp Med Biol 1239, 21–40 (2020). 10.1007/978-3-030-38062-5_3

36 Belyantseva, I. A. et al. Myosin-XVa is required for tip localization of whirlin and differential elongation of hair-cell stereocilia. Nature cell biology 7, 148–156 (2005). 10.1038/ncb1219

37 Manor, U. et al. Regulation of stereocilia length by myosin XVa and whirlin depends on the actin-regulatory protein Eps8. Curr Biol 21, 167–172 (2011). 10.1016/j.cub.2010.12.046

38 Moreland, Z. G. et al. Myosin-driven Nucleation of Actin Filaments Drives Stereocilia Development Critical for Hearing. bioRxiv, 2021.2007.2009.451618 (2021). 10.1101/2021.07.09.451618

39 Liu, H. et al. Myosin III-mediated cross-linking and stimulation of actin bundling activity of Espin. Elife 5 (2016). 10.7554/eLife.12856

40 Salles, F. T. et al. Myosin IIIa boosts elongation of stereocilia by transporting espin 1 to the plus ends of actin filaments. Nature cell biology 11, 443–450 (2009). 10.1038/ncb1851

41 Lelli, A. et al. Class III myosins shape the auditory hair bundles by limiting microvilli and stereocilia growth. The Journal of cell biology 212, 231–244 (2016). 10.1083/jcb.201509017

42 Grillet, N. et al. Harmonin mutations cause mechanotransduction defects in cochlear hair cells. Neuron 62, 375–387 (2009). 10.1016/j.neuron.2009.04.006

43 Grati, M. & Kachar, B. Myosin VIIa and sans localization at stereocilia upper tip-link density implicates these Usher syndrome proteins in mechanotransduction. Proceedings of the National Academy of Sciences of the United States of America 108, 11476–11481 (2011). 10.1073/pnas.1104161108

44 Kazmierczak, P. et al. Cadherin 23 and protocadherin 15 interact to form tip-link filaments in sensory hair cells. Nature 449, 87–91 (2007). 10.1038/nature06091

45 Lefevre, G. et al. A core cochlear phenotype in USH1 mouse mutants implicates fibrous links of the hair bundle in its cohesion, orientation and differential growth. Development 135, 1427–1437 (2008). 10.1242/dev.012922

46 Belyantseva, I. A., Boger, E. T. & Friedman, T. B. Myosin XVa localizes to the tips of inner ear sensory cell stereocilia and is essential for staircase formation of the hair bundle. Proceedings of the National Academy of Sciences of the United States of America 100, 13958–13963 (2003). 10.1073/pnas.2334417100

47 Umeki, N. et al. The tail binds to the head-neck domain, inhibiting ATPase activity of myosin VIIA. Proceedings of the National Academy of Sciences of the United States of America 106, 8483–8488 (2009). 10.1073/pnas.0812930106

48 Yang, Y. et al. A FERM domain autoregulates Drosophila myosin 7a activity. Proceedings of the National Academy of Sciences of the United States of America 106, 4189–4194 (2009). 10.1073/pnas.0808682106

49 Zhuravlev, P. I., Lan, Y., Minakova, M. S. & Papoian, G. A. Theory of active transport in filopodia and stereocilia. Proceedings of the National Academy of Sciences of the United States of America 109, 10849–10854 (2012). 10.1073/pnas.1200160109

50 Li, A., Xue, J. & Peterson, E. H. Architecture of the mouse utricle: macular organization and hair bundle heights. Journal of neurophysiology 99, 718–733 (2008). 10.1152/jn.00831.2007

51 Belyantseva, I. A. Helios® Gene Gun-Mediated Transfection of the Inner Ear Sensory Epithelium: Recent Updates. Methods Mol Biol 1427, 3–26 (2016). 10.1007/978-1-4939-3615-1_1

52 Grimm, J. B. et al. A General Method to Improve Fluorophores Using Deuterated Auxochromes. JACS Au 1, 690–696 (2021). 10.1021/jacsau.1c00006

53 Sakai, T., Umeki, N., Ikebe, R. & Ikebe, M. Cargo binding activates myosin VIIA motor function in cells. Proceedings of the National Academy of Sciences of the United States of America 108, 7028–7033 (2011). 10.1073/pnas.1009188108

54 Szent-Gyorgyi, A. G. Meromyosins, the subunits of myosin. Archives of biochemistry and biophysics 42, 305–320 (1953). 10.1016/0003-9861(53)90360-9

55 Lowey, S. Myosin Substructure: Isolation of a Helical Subunit from Heavy Meromyosin. Science 145, 597–599 (1964). 10.1126/science.145.3632.597

56 Matoo, S. et al. Comparative analysis of the MyTH4-FERM myosins reveals insights into the determinants of actin track selection in polarized epithelia. Molecular biology of the cell 32, ar30 (2021). 10.1091/mbc.E20-07-0494

57 Clackson, T. et al. Redesigning an FKBP-ligand interface to generate chemical dimerizers with novel specificity. Proceedings of the National Academy of Sciences of the United States of America 95, 10437–10442 (1998). 10.1073/pnas.95.18.10437

58 Sato, O. et al. Human myosin VIIa is a very slow processive motor protein on various cellular actin structures. The Journal of biological chemistry 292, 10950–10960 (2017). 10.1074/jbc.M116.765966

59 Kerber, M. L. et al. A novel form of motility in filopodia revealed by imaging myosin-X at the single-molecule level. Curr Biol 19, 967–973 (2009). 10.1016/j.cub.2009.03.067

60 Spink, B. J., Sivaramakrishnan, S., Lipfert, J., Doniach, S. & Spudich, J. A. Long single alpha-helical tail domains bridge the gap between structure and function of myosin VI. Nat Struct Mol Biol 15, 591–597 (2008). 10.1038/nsmb.1429

61 Thirumurugan, K., Sakamoto, T., Hammer, J. A., 3rd, Sellers, J. R. & Knight, P. J. The cargo-binding domain regulates structure and activity of myosin 5. Nature 442, 212–215 (2006). 10.1038/nature04865

62 Boeda, B. et al. Myosin VIIa, harmonin and cadherin 23, three Usher I gene products that cooperate to shape the sensory hair cell bundle. EMBO J 21, 6689–6699 (2002). 10.1093/emboj/cdf689

63 Siemens, J. et al. The Usher syndrome proteins cadherin 23 and harmonin form a complex by means of PDZ-domain interactions. Proceedings of the National Academy of Sciences of the United States of America 99, 14946–14951 (2002). 10.1073/pnas.232579599

64 Adato, A. et al. Interactions in the network of Usher syndrome type 1 proteins. Hum Mol Genet 14, 347–356 (2005). 10.1093/hmg/ddi031

65 Senften, M. et al. Physical and functional interaction between protocadherin 15 and myosin VIIa in mechanosensory hair cells. The Journal of neuroscience : the official journal of the Society for Neuroscience 26, 2060–2071 (2006). 10.1523/JNEUROSCI.4251-05.2006

66 Fitz, G. N., Weck, M. L., Bodnya, C., Perkins, O. L. & Tyska, M. J. Protrusion growth driven by myosin-generated force. Developmental cell 58, 18–33 e16 (2023). 10.1016/j.devcel.2022.12.001

67 Grati, M. et al. MYO3A Causes Human Dominant Deafness and Interacts with Protocadherin 15-CD2 Isoform. Hum Mutat 37, 481–487 (2016). 10.1002/humu.22961

68 Inobe, T. & Nukina, N. Rapamycin-induced oligomer formation system of FRB-FKBP fusion proteins. J Biosci Bioeng 122, 40–46 (2016). 10.1016/j.jbiosc.2015.12.004

69 Raval, M. H. et al. Impact of the Motor and Tail Domains of Class III Myosins on Regulating the Formation and Elongation of Actin Protrusions. The Journal of biological chemistry 291, 22781–22792 (2016). 10.1074/jbc.M116.733741

70 Rayment, I. et al. Three-dimensional structure of myosin subfragment-1: a molecular motor. Science 261, 50–58 (1993). 10.1126/science.8316857

71 Castillo, M., Freire, E. & Romero, V. I. Primary ciliary dyskinesia diagnosis and management and its implications in America: a mini review. Front Pediatr 11, 1091173 (2023). 10.3389/fped.2023.1091173

72 Hildebrandt, F., Benzing, T. & Katsanis, N. Ciliopathies. The New England journal of medicine 364, 1533–1543 (2011). 10.1056/NEJMra1010172

73 VanDussen, K. L. et al. Abnormal Small Intestinal Epithelial Microvilli in Patients With Crohn’s Disease. Gastroenterology 155, 815–828 (2018). 10.1053/j.gastro.2018.05.028

74 Modl, B. et al. Defects in microvillus crosslinking sensitize to colitis and inflammatory bowel disease. EMBO Rep 24, e57084 (2023). 10.15252/embr.202357084

75 Jacquemet, G., Hamidi, H. & Ivaska, J. Filopodia in cell adhesion, 3D migration and cancer cell invasion. Current opinion in cell biology 36, 23–31 (2015). 10.1016/j.ceb.2015.06.007

76 Krey, J. F. & Barr-Gillespie, P. G. Molecular Composition of Vestibular Hair Bundles. Cold Spring Harb Perspect Med 9 (2019). 10.1101/cshperspect.a033209

77 Stauffer, E. A. et al. Fast adaptation in vestibular hair cells requires myosin-1c activity. Neuron 47, 541–553 (2005). 10.1016/j.neuron.2005.07.024

78 Self, T. et al. Role of myosin VI in the differentiation of cochlear hair cells. Developmental biology 214, 331–341 (1999). 10.1006/dbio.1999.9424

79 Ahmed, Z. M. et al. Mutations of MYO6 are associated with recessive deafness, DFNB37. Am J Hum Genet 72, 1315–1322 (2003). 10.1086/375122

80 Melchionda, S. et al. MYO6, the human homologue of the gene responsible for deafness in Snell’s waltzer mice, is mutated in autosomal dominant nonsyndromic hearing loss. Am J Hum Genet 69, 635–640 (2001). 10.1086/323156

81 Liu, X. Z. et al. Mutations in the myosin VIIA gene cause non-syndromic recessive deafness. Nat Genet 16, 188–190 (1997). 10.1038/ng0697-188

82 Weil, D. et al. The autosomal recessive isolated deafness, DFNB2, and the Usher 1B syndrome are allelic defects of the myosin-VIIA gene. Nat Genet 16, 191-193 (1997). 10.1038/ng0697-191

83 Dionne, G. et al. Mechanotransduction by PCDH15 Relies on a Novel cis-Dimeric Architecture. Neuron 99, 480–492 e485 (2018). 10.1016/j.neuron.2018.07.006

84 Jaiganesh, A. et al. Zooming in on Cadherin-23: Structural Diversity and Potential Mechanisms of Inherited Deafness. Structure 26, 1210–1225 e1214 (2018). 10.1016/j.str.2018.06.003

85 Verpy, E. et al. A defect in harmonin, a PDZ domain-containing protein expressed in the inner ear sensory hair cells, underlies Usher syndrome type 1C. Nat Genet 26, 51–55 (2000). 10.1038/79171

86 Bahloul, A. et al. Cadherin-23, myosin VIIa and harmonin, encoded by Usher syndrome type I genes, form a ternary complex and interact with membrane phospholipids. Hum Mol Genet 19, 3557–3565 (2010). 10.1093/hmg/ddq271

87 Caberlotto, E. et al. Usher type 1G protein sans is a critical component of the tip-link complex, a structure controlling actin polymerization in stereocilia. Proceedings of the National Academy of Sciences of the United States of America 108, 5825–5830 (2011). 10.1073/pnas.1017114108

88 Indzhykulian, A. A. et al. Molecular remodeling of tip links underlies mechanosensory regeneration in auditory hair cells. PLoS Biol 11, e1001583 (2013). 10.1371/journal.pbio.1001583

89 Elferich, J. et al. Molecular structures and conformations of protocadherin-15 and its complexes on stereocilia elucidated by cryo-electron tomography. Elife 10 (2021). 10.7554/eLife.74512

90 De La Cruz, E. M. & Ostap, E. M. Relating biochemistry and function in the myosin superfamily. Current opinion in cell biology 16, 61–67 (2004). 10.1016/j.ceb.2003.11.011

91 Les Erickson, F., Corsa, A. C., Dose, A. C. & Burnside, B. Localization of a class III myosin to filopodia tips in transfected HeLa cells requires an actin-binding site in its tail domain. Molecular biology of the cell 14, 4173–4180 (2003). 10.1091/mbc.e02-10-0656

92 Quintero, O. A. et al. Intermolecular autophosphorylation regulates myosin IIIa activity and localization in parallel actin bundles. The Journal of biological chemistry 285, 35770–35782 (2010). 10.1074/jbc.M110.144360

93 Houdusse, A. & Titus, M. A. The many roles of myosins in filopodia, microvilli and stereocilia. Curr Biol 31, R586–R602 (2021). 10.1016/j.cub.2021.04.005

94 Miyoshi, T. et al. Actin turnover-dependent fast dissociation of capping protein in the dendritic nucleation actin network: evidence of frequent filament severing. The Journal of cell biology 175, 947–955 (2006). 10.1083/jcb.200604176

95 Narayanan, P. et al. Length regulation of mechanosensitive stereocilia depends on very slow actin dynamics and filament-severing proteins. Nat Commun 6, 6855 (2015). 10.1038/ncomms7855

96 Jia, S., Yang, S., Guo, W. & He, D. Z. Fate of mammalian cochlear hair cells and stereocilia after loss of the stereocilia. The Journal of neuroscience : the official journal of the Society for Neuroscience 29, 15277–15285 (2009). 10.1523/JNEUROSCI.3231-09.2009

97 Yildiz, A. et al. Myosin V walks hand-over-hand: single fluorophore imaging with 1.5-nm localization. Science 300, 2061–2065 (2003). 10.1126/science.1084398

98 Wu, Y. et al. Reflective imaging improves spatiotemporal resolution and collection efficiency in light sheet microscopy. Nat Commun 8, 1452 (2017). 10.1038/s41467-017-01250-8

99 Endesfelder, U. & Heilemann, M. Direct stochastic optical reconstruction microscopy (dSTORM). Methods Mol Biol 1251, 263–276 (2015). 10.1007/978-1-4939-2080-8_14

100 Banaz, N., Makela, J. & Uphoff, S. Choosing the right label for single-molecule tracking in live bacteria: side-by-side comparison of photoactivatable fluorescent protein and Halo tag dyes. J Phys D Appl Phys 52, 064002 (2019). 10.1088/1361-6463/aaf255

101 Lamptey, R. N. L. et al. A Review of the Common Neurodegenerative Disorders: Current Therapeutic Approaches and the Potential Role of Nanotherapeutics. Int J Mol Sci 23 (2022). 10.3390/ijms23031851

102 Karginov, A. V., Ding, F., Kota, P., Dokholyan, N. V. & Hahn, K. M. Engineered allosteric activation of kinases in living cells. Nat Biotechnol 28, 743–747 (2010). 10.1038/nbt.1639

103 Sun, X., Tie, H. C., Chen, B. & Lu, L. Glycans function as a Golgi export signal to promote the constitutive exocytic trafficking. The Journal of biological chemistry 295, 14750–14762 (2020). 10.1074/jbc.RA120.014476

104 Edelstein, A. D. et al. Advanced methods of microscope control using µManager software. J Biol Methods 1 (2014). 10.14440/jbm.2014.36

105 Schneider, C. A., Rasband, W. S. & Eliceiri, K. W. NIH Image to ImageJ: 25 years of image analysis. Nature methods 9, 671–675 (2012). 10.1038/nmeth.2089

106 Ardiel, E. L. et al. Visualizing Calcium Flux in Freely Moving Nematode Embryos. Biophysical journal 112, 1975–1983 (2017). 10.1016/j.bpj.2017.02.035

107 Nagashima, S., Aoki, T., Higuchi, T. & Kobayashi, K. in 2006 International Symposium on Intelligent Signal Processing and Communications. 701–704.

108 Guo, M. et al. Accelerating iterative deconvolution and multiview fusion by orders of magnitude. (2019). 10.1101/647370

109 Guo, M. et al. Rapid image deconvolution and multiview fusion for optical microscopy. Nat Biotechnol 38, 1337–1346 (2020). 10.1038/s41587-020-0560-x

110. Schindelin, J., et al. Fiji: an open-source platform for biological-image analysis. Nature methods 9, 676-682 (2012). 10.1038/nmeth.2019

111 Sakai, T. et al. Structure and Regulation of the Movement of Human Myosin VIIA. The Journal of biological chemistry 290, 17587–17598 (2015). 10.1074/jbc.M114.599365

112 Wu, L., Pan, L., Wei, Z. & Zhang, M. Structure of MyTH4-FERM domains in myosin VIIa tail bound to cargo. Science 331, 757–760 (2011). 10.1126/science.1198848

113 Liu, R. et al. A binding protein regulates myosin-7a dimerization and actin bundle assembly. Nat Commun 12, 563 (2021). 10.1038/s41467-020-20864-z

